# Genetic and functional odorant receptor variation in the Homo lineage

**DOI:** 10.1101/2021.09.13.460146

**Authors:** Claire A. de March, Hiroaki Matsunami, Masashi Abe, Matthew Cobb, Kara C. Hoover

**Author notes:** These authors contributed equally to this work” (Claire A. de March, Kara C. Hoover).

## Abstract

Using ancient DNA sequences, we explored the function of olfactory receptor genes in the genus Homo. Humans, Neandertals, and Denisovans independently adapted to a wide range of geographic environments and the odours produced by their food. Variations in their odorant receptor protein sequence and structure resulted in variation in detection and perception. Studying thirty olfactory receptor genes, we found our relatives showed highly conserved receptor structures, but Homo sapiens did not. Variants led to changes in sensitivity to some odors, but no change in specificity, indicating a common olfactory repertoire in our genus. Diversity of geographic adaptations in H. sapiens may have produced greater functional variation in our lineage, increasing our olfactory repertoire and expanding our adaptive capacity.

**One-Sentence Summary:** Using ancient DNA we studied the sense of smell in our extinct ancestors and in our relatives, Denisovans and Neanderthals

## Main Text

Terrestrial animals smell by binding odorant molecules to odorant receptors (ORs) but there is limited knowledge of odorant-OR and genotype-phenotype associations. Variation in mammalian ORs is strongly linked to ecological and dietary niche (1, 2). The human genus Homo underwent the most radical ecological niche expansion of all primates when migrating out of Africa and adapting to diverse global environments (3). Denisovans and Neandertals ancestors dispersed from Africa earlier than present-day humans (4) (∼750,000 versus 65,000 years ago) and separated from each other ∼300,000 years ago (5) (Fig. S1). Neandertals were geographically wide-ranging (western Europe, Middle East, Asia) while Denisovans were geographically constrained to Siberia (6, 7), the Tibetan plateau (8), and possibly beyond Wallace’s Line (9). Olfactory stimuli from divergent environments following independent dispersals from Africa may have left traces of variation in Homo ORs. Among present-day humans, changes in OR function are linked to major evolutionary dietary shifts, such as scavenging, hunting, animal milk consumption, cooking, domestication (1, 10–14). But, what about the gap between Homo migrations and human-specific changes in ORs?

Using published ancient DNA sequences to analyze genetic variation for population structure and test functional differences in gene variants for ecological differentiation, we assess whether there is a shared Homo olfactory repertoire (range of detectable odors). We previously explored genetic and functional variation in present-day humans, Altai Neandertal, and Denisovan for OR7D4 (15). We extended our study to 29 additional open reading frames (Table S1) for ORs with known human receptor-odor (16, 17) and two additional Neandertals (Chagyrskaya, Vindija) and one ancient human (Ust’-Ishim) who lived in the same Altai montane locality (Table S2). We used 1000 Genomes for present-day humans (Table S3) (6). We found that novel variants alter sensitivity but not specificity of OR function and conclude that the olfactory repertoires of extinct lineages were highly overlapping. We also argue that expansion of the human olfactory repertoire occurred after our split with other migratory members of our genus.

## Results

### Genetic variation

Extinct lineages and Ust’-Ishim had fewer DNA and protein variants than 1000 Genomes (Fig. 1, Table S4)—on average 0.19% of nucleotides across 17 genes (or 0.11% when dividing across all 30 genes, including those matching the human wild type) compared to 0.82% of nucleotides across 30 genes in 1000 Genomes. Ancient populations were more prone to genetic drift (fixation and loss) due to smaller effective population sizes and living in small, isolated communities (18). Extinct lineages exhibit a pattern of OR gene conservation and present-day humans exhibit a pattern of tolerating variation.

**Fig. 1.**
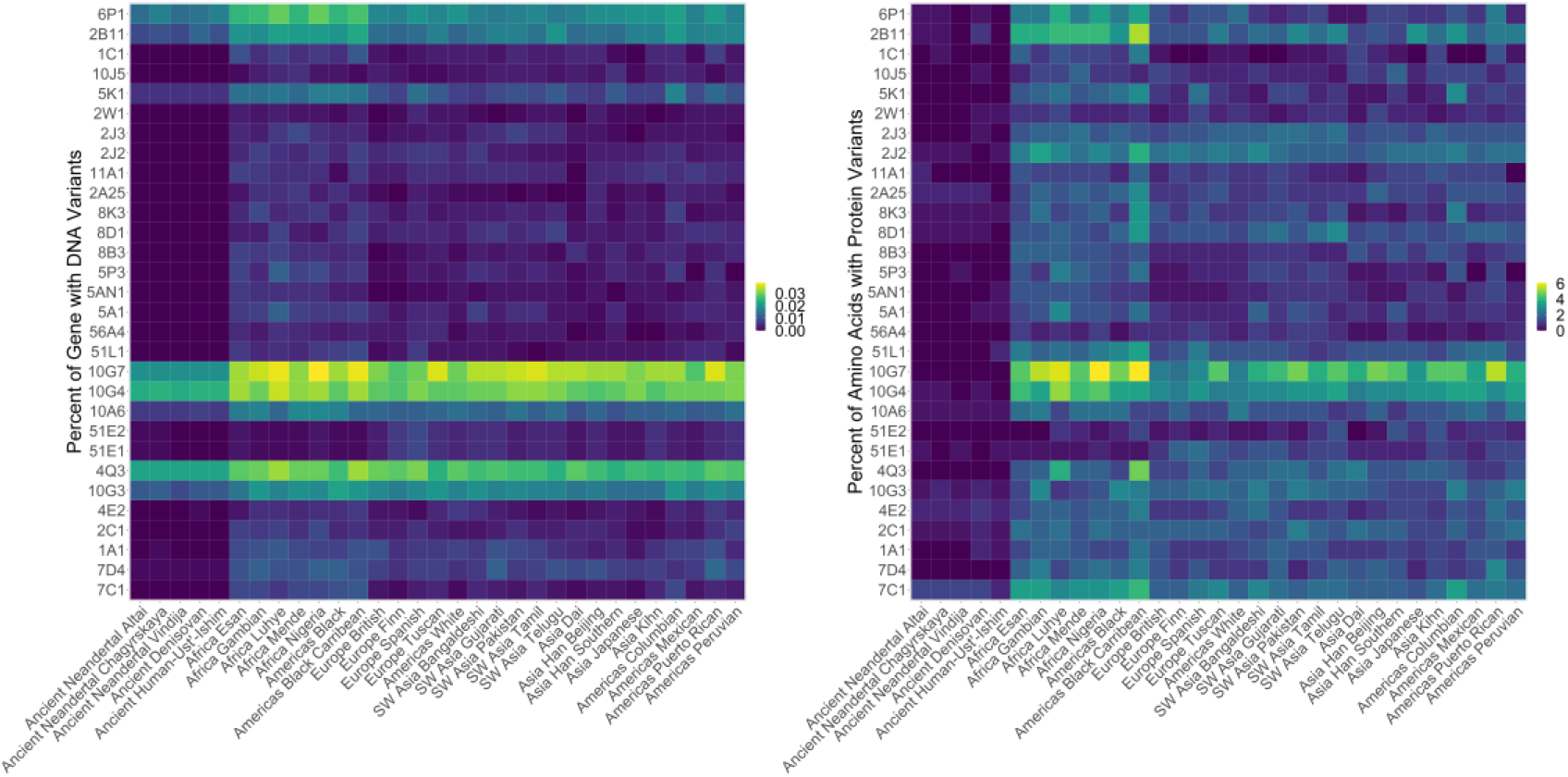
Percent OR variation. Total variant count per gene divided by total basepairs in gene for each population and based on raw counts for ancient populations and raw count for the consensus sequence of each gene for each of 26 groups in 1000 Genomes.

The fixation index (Fst) measures genetic variance due to population structure (typically weighted by population size) and ranges from 0 (no differentiation) to 1 (complete differentiation) (19). The Fst values by gene for 1000 Genomes populations are lower than other large-bodied mammals with wide geographic dispersal (Table S5) (20). OR5P3 fixation was the highest at 11.4%, the lower limit of possible significant differentiation by population structure. The 1000 Genomes populations are not highly differentiation based on the 30 ORs used in this study an Fst mean of 4%. In contrast, the genus Homo (1000 Genomes and ancient samples) has an Fst mean of 11%, the lower end of possible significant differentiation by population structure. There was variation among Fst by gene, however. Ten Fst values were <1% (OR2W1, OR2J2, OR4Q3, OR5K1, OR6P1, OR7D4, OR8B3, OR10G7, OR11A1, OR51L1), which suggests the sort of random distribution associated with panmixia or genetic conservation. Of the 13 genes with Fst > 12%, seven were >20% (1C1, 5AN1, 7C1, 10G3, 10J5, 51E1, 51E2), which suggests the sort of distribution associated with high population differentiation. Of the genes with high Fst, only two had novel variants in extinct populations, which suggests that the small number of novel variants are not highly influential in differentiating populations (Table S7). The greater number of genes with high Fst indicate that Homo used to be more structured by population than present-day humans.

Looking at the distribution of variants, two ancient genes were identical to the human wild type (OR4Q3, OR8B3) and twenty contained variants also observed with 1000 Genomes—shared variation (Table S6), suggesting these variants occurred prior to global divergence of Homo.

Only 11 genes, with a total of 14 variants, were unique to ancient samples (not found in 1000 Genomes). These genes may well be ones that were influenced by olfactory stimuli in divergent environments following dispersals from Africa. Denisovan had nine novel variants (of which two were synonymous) compared to the Neandertal five (of which 2 were synonymous). No novel variants were observed in the ancient human Ust’-Ishim.

Amino acid sequences for the ORs studied were used to form a cladogram to explore the relationships of our samples (Fig. S2). Extinct lineages formed a clade with Vindija Neandertal the most distinct, which was unexpected because genome-wide studies have indicated that Vindija is most genetically similar to Chagyrskaya Neandertal (18). The extinct clade was closest to the ancient human Ust’-Ishim and then to East and South Asian present-day humans—the latter groups harbor genetic signatures of introgression with Neandertals and Denisovans (9).

### Functional Variation

Because gene function in not reliably predictable for ORs from sequence data (21, 22), we directly measured the functional responses of ORs containing novel variants.

Each OR protein, expressed in a cell line, was screened against seven odorants previously identified in the literature as evoking responses: OR1A1 (21, 23), OR1C1 (22), OR2C1 (21), OR10J5 (21), OR5P3 (21), and OR10G3 (22). Dose response assays for the top screening responses included seven concentrations of the odors delivered separately.

There were only three Neandertal genes containing novel variants. Their dose responses were not correlated with those of present-day humans (Figs. 2A, 2C, 2D). Only 1C1 had a detectable response but significantly lower than that of present-day humans (Figs. 3A, 3C, 3D, 4, S2, S3).

**Fig. 2.**
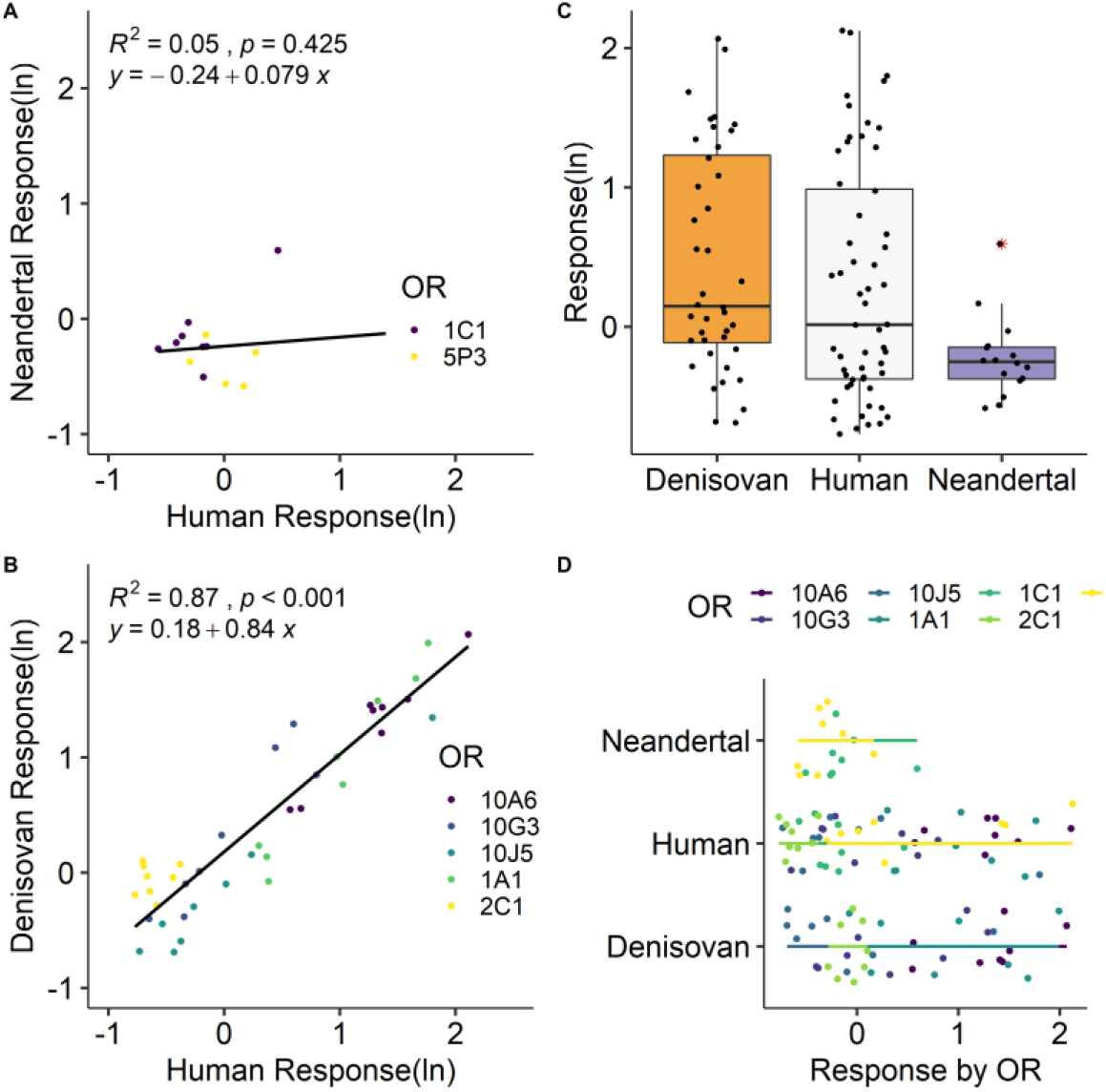
Regression results for dose responses. **A**. Human and Denisovan, **B**. Human and Neandertal. **C**. Boxplots of dose responses for all samples showing median, box boundaries (first and third quartiles), and two whiskers (upper whisker extends to the largest value no further than 1.5 inter-quartile range from third quartile; lower whisker extends to the smallest value at most 1.5 inter-quartile range of first quartile, outliers identified with red asterisk), **D**. Activity Indesx by OR for all three samples.

**Fig. 3:**
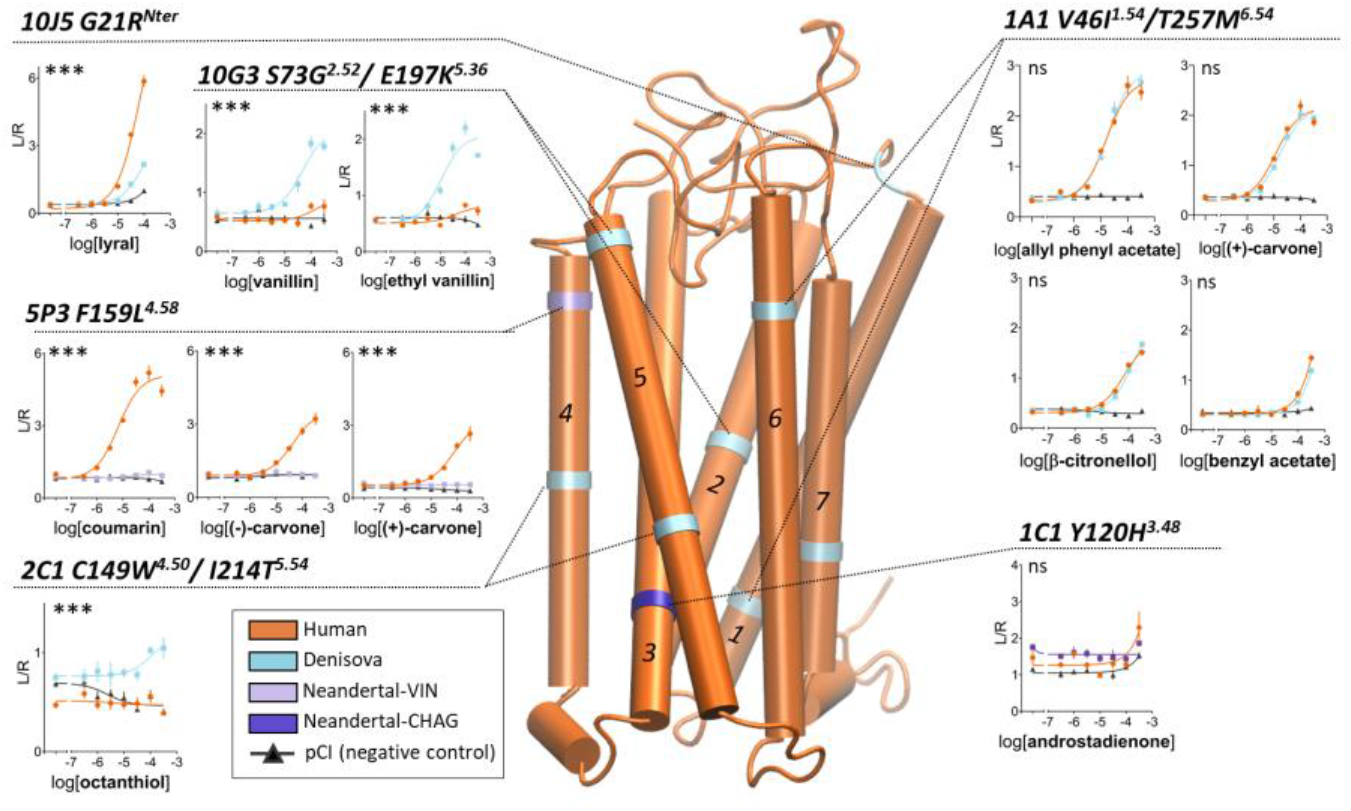
Homology model of the human consensus odorant receptor. To illustrate location of variants, panels show dose-response for odorants that were significantly activating ORs in screening. (FigS3; 10G3 and 2C1 both have one shared and one novel mutation). The x-axis of panels represents the odorant concentration (M) and the y-axis the normalized luminescence generated by the activated OR. Error bars are SEM (standard error of the mean).

Despite the higher number of novel OR variants in the Denisovans and higher dose responses compared to present-day humans (Figs. 2C, 2D), the OR responses for six genes and those for human reference were significantly correlated (R2 = 0.87) (Fig. 2B). Tested Denisovan ORs were less sensitive to odors that present-day humans perceive as floral but much more sensitive to odors perceived as spicy, balsamic, or unpleasant (e.g., sulfur 4x greater and balsamic 3x greater than in present-day humans) (Table 1). Higher dose responses in Denisovan ORs appear to be driven by two amino acid variations in two of the ORs with novel variants (Fig. 3, Fig. S3).

**Table 1.**
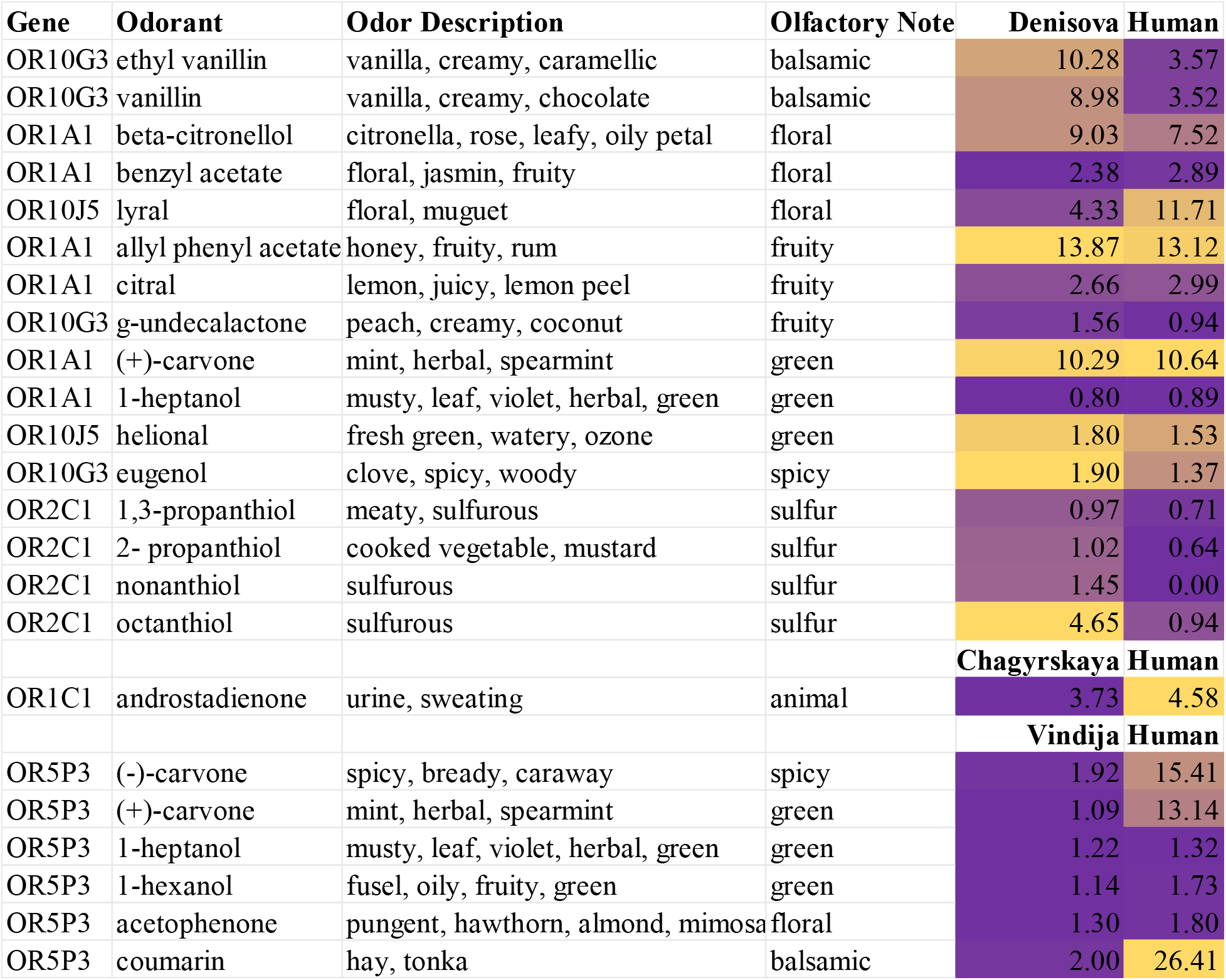
Comparison of OR activity index for human and extinct lineages. Color coding is from low (purple) to high (mustard).

Despite using the same DNA sequence as previous studies of 2B11 (22) and 6P1 (17), the human reference for both did not respond to any of the ∼350 odors (100µM) against which they were tested (Fig. S4)—neither did the extinct linages. In previous studies, OR2B11 (22) and OR6P1 (17) responded strongly to coumarin and anisaldehyde, respectively, at concentrations higher than 100µM. We found that such concentrations often cause cell toxicity or OR non-specific cell responses.

OR1A1. All but 2-heptanone induced a response in the screening assay (Fig. S3A) and the highest responses were for honey (particularly for Denisovan) (Table 1, Fig. 3). Neither of the two Denisovan variants (V461I1.54, T257M6.54) were in an amino acid region critical for mammalian OR function nor were they involved in the odorant binding cavity—perhaps explaining their minimal functional impact.

OR1C1. The only significant response was from Chagyrskaya Neandertal to androstadienone (Fig. S3B) and it was weak (Table 1; Fig. 3). Chagyrskaya Neandertal 1C1 variant Y120H3.48 is part of the highly conserved MAY3.48DRY motif involved in the activation of mammalian ORs (Fig. 3, Fig. S3B), which might explain why this variant alters function.

OR2C1. Screening assay responses were strong but not statistically significant (Fig. S3A). The dose response assay for octanethiol produced a statistically significant response in the Denisovan version of this OR (Table 1, Fig. 3). The shared C149W4.50 corresponds to the conserved W4.50. The W residue is highly conserved in GPCRs but less so in ORs (58%). The location of the novel Denisovan I214T5.54 in TM5 is below residues involved in canonical ligand binding cavity and it points into the receptor rather than the surface. In addition, prior functional tests for C149 found a similar response (17). The C149W allele may stop protein function and may have produced octanethiol-specific anosmia.

OR5P3. In the screening assay, Vindija Neandertal did not have a significant response but the human reference responded to five of the seven odorants (Fig. S3B). Vindija dose responses, which included higher concentrations of the top three responses for present-day humans (courmarin and both enantiomers of carvone), did not exceed control (Fig. 3). The cell surface expression for Vindija indicated that the OR proteins was present at the cell surface, albeit at a slightly lower level than that in present-day humans. Vindija F159L4.58 is in the extracellular part of TM4 (Fig. 3), near 4.53, which is involved in mouse OR trafficking (24). We observed slightly lower trafficking of the Vindija protein. A similar mutation (S155A4.56) in human OR1A2 decreases in vitro responses to (S)-(-)-citronellal (23). If OR5P3 F159L4.58 is involved in odorant binding, the mutation of this position from phenylalanine to leucine might prevent the π-π stacking interaction between the aromatic residue and coumarin. We conclude that the Vindija protein is not functional and this might be attributable to many potential reasons (a few examples are lack of odorant binding, or fail in activation mechanism, or fail to bind the G protein).

OR10G3. The screening assay revealed significant responses for all seven odors in the Denisovan OR and in the human reference (Fig. S3A). Denisovan variants had significantly stronger dose responses to vanillin and ethyl vanillin compared to present-day humans (Table 1; Fig. 3). Neither of the Denisovan variants ( S73G2.52, E197K5.36) were located in conserved amino acid regions (Fig. 3). TM2 is not involved in odorant binding or receptor function, implying that S73G2.52 probably did not change the receptor response. TM5 (E197K5.36) forms part of the binding cavity but position 5.36 is located at the very limit of ECL2. K at this position is a rare residue in present-day human ORs (3.6%), suggesting a functional, adaptative reason for this change. The location of the variant suggests it may be involved in ligand entry.

OR10J5. There were three significant screening responses to lyral and helional and eugenol (Fig. S3A). The Denisovan response to top odor lyral was lower than that of the human reference (Table 1, Fig. 3). The G21RNter variant of 10J5 found in the Denisovan is located at the very end of the N terminal end, just before the start of TM1. The role of this region in OR function is undetermined.

## Discussion

Ten novel missense variants were located in 8 genes (out of 30), with five being functionally different from present-day humans (1C1, 2C1, 5P3, 10G3, 10J5), one the same (1A1) and two without identifiable ligands (2B11, 6P1). Given the small percentage of genes with variants altering OR function, members of the genus Homo likely shared an olfactory repertoire, with Neandertals and Denisovans smelling the same range of odors we do but having different dose responses to those odors. When OR function was altered by a novel OR, the difference was in sensitivity rather than specificity. Novel Denisovan OR variants (1A1, 2C1, 10G3, 10J5) were twice as responsive as human equivalents to odors present-day humans perceive as spicy, balsamic, and unpleasant (Table 1), but not to odors perceived as floral. Novel Neandertal variants were three times less responsive than human ORs, including reduced responses to odors perceived as green, floral, and spicy (Table 1). There is some correlation in Neandertal skull morphology that suggests their olfactory bulbs were smaller than present-day humans (6), but the link between bulb size and olfactory acuity is unclear (25, 26).

The Denisovan ORs strong response to honey may be ecologically significant. Honey is the most energy-dense natural food and is a prized component of living hunter-gatherer diets (except where bees are rare or absent)—even great apes have a ‘honey tooth’ (27). Energy-dense foods like sugars are sought by larger-brained primates (28) and, based on oral microbiome data, Neandertals and present-day humans share functional adaptations in nutrient metabolism including starch digestion that are not found in our closest ape relatives (29). The high response to vanilla odors suggests a potential response to sweet things—an odor-taste pairing common in present-day humans (30).

Local ecological adaptive pressures may have acted on ORs in extinct lineages to produce the few novel variants observed, but extinct lineages were less variable in OR genes and proteins compared to 1000 Genomes. While differences in sample sizes might account for some of the striking differences, the reduced variation is probably due to genetic drift effect or conservation. Small effective population sizes cause bottlenecks and small, isolated populations (18) are prone to genetic drift. Purifying selection has been observed in chimpanzee ORs compared to a mix of relaxed and positive selection in human ORs (31) and extinct lineages may also have been subject to this, as evidenced by fewer variants that mostly code for synonymous proteins (compared to protein variation in 1000 Genomes). The mean of Fst values across genes comparing 1000 Genomes to extinct lineaages (11%) is higher than those for 1000 Genomes population comparisons (4%), which suggests that there are structural differences between them and us that reflect both explanations—drift and conservation. Based on our data the last common ancestor shared by Homo (present-day humans, Neandertals, Denisova, and others) and Pan (chimpanzees, bonobos) had a conserved set of ORs. Present-day humans derived away from the pattern of conservation more recently, with evolutionary pressure toward increased missense variation.

Despite having a shared repertoire with Neandertals and Denisovans, present-day humans are novel in their highly variable OR repertoire which may reflect cultural adaptations following migrations from Africa. Relaxed selection on OR genes for groups no longer engaging in traditional lifestyles is possible (32). 1000 Genomes groups outside Africa are less variable in most OR genes than those in Africa and OR gene enrichment is observed in African hunter-gatherer groups (Hadza and Pygmies) but not African agricultural (Yoruba) and pastoral (Maasai) groups (33). Tanzanian Sandawe hunter-gatherers show no OR allelic enrichment, however, which undermines the case for relaxed selection (33). High allelic diversity and OR generalization on a broad scale may have functional implications, such as increasing the effective size of the olfactory repertoire (34). Understanding our unique OR allelic diversity is an important challenge.

Our data provide insights into how the dispersal of human lineages outside of Africa (Denisova, Neandertal, ancient human) may have affected olfactory gene repertoire and function.

Understanding the evolutionary genetics and functional significance of observed OR allelic variability in and almong human populations and extinct relatives sheds light on the role of olfaction in key aspects of human culture, and perhaps our current success as a global species.

## Acknowledgments

The high-performance computing and data storage resources operated by the Research Computing Systems Group at the University of Alaska Fairbanks, Geophysical Institute.

## Funding

Provide complete funding information, including grant numbers, complete funding agency names, and recipient’s initials. Each funding source should be listed in a separate paragraph.

US National Science Foundation Award 1550409 (KCH) National Institutes of Health grant K99DC018333 (CADM) US National Science Foundation Award 1556207 (HM) National Institutes of Health grant DC014423 (HM) National Institutes of Health grant DC016224 (HM)

## Author contributions

Conceptualization: KCH, HM, MC

Methodology: KCH, CADM, HM, MC

Investigation: KCH, CADM, MA

Visualization: KCH, CADM

Funding acquisition: KCH, CADM, HM

Project administration: KCH, HM

Supervision: KCH, HM

Writing – original draft: KCH, CADM

Writing – review & editing: KCH, CADM, MC, HM

## Competing interests

Authors declare that they have no competing interests.

## Data and materials availability

Data Availability. Raw data for this project and data derived from it are available in at https://github.com/kchoover14/OldNoses and are usable under the license provided in the repository.

### Code Availability

Code for VCF data scraping and code for analysis in Tables S4-S5 and Figs. 1-2 and Figs. S1-S2, S7-S9 are available at https://github.com/kchoover14/OldNoses and are usable under the license provided in the repository.

## Supplementary Materials

### Materials and Methods

#### Materials

We cataloged variants in high quality paleogenomic sequence data produced by the Max Planck Institute Leipzig (Table S2) for Neandertals (Altai, Chagyrskaya, Mezmaiskaya), Denisovan 3 (the only high quality Denisovan genome, finger phalanx), and an ancient human hunter-gatherer from Siberia (Ust’-Ishim). We used the data generated by snpAD in /neandertal/vindija/vcf, an ancient DNA damage-aware genotyper.1 The VCF reference genome for genomes analyzed was hg19/GRCh37. Only variants with a minimum genotype quality of 20 (GC20) were used for downstream analysis. Our sample of present-day humans was from the 1000 Genomes dataset (35) consisting of over 2,500 individuals from 26 populations in Africa, the Americas, Europe, and Asia (Table S3). The 26 groups in 1000 Genomes were comprised of individuals with at least 3 out of 4 grandparents identifying membership in the group. While there are some groups engaging in pastoral or traditional horticulture/farming lifestyles, most practice mixed subsistence economies and lead western lifestyles. Although the draft genomes of Neandertals and Denisovans were reported as having more nonfunctional ORs than present-day humans (36), these genomes have since been revised (37, 38). The bioinformatic identification of olfactory pseudogenes has been experimentally challenged (39) by data showing the receptors containing coding sequence regions split into separate exons are conserved across mammals and expressed at the same level as protein-coding receptors with a single exon (40).

We focused on 30 OR genes shown to generate functional response data in previous studies (16) (Table S1). Gene regions were targeted using NCBI and RefSeq ranges—in the case of eight genes that contained up- and downstream sequences, the region was cut to the protein coding portion of the gene to be consistent with the other 22 genes that contained no up- or downstream areas.

### Methods

#### Variant calling

Max Planck and 1000 Genomes Project both used the reference genome, hg19/GRCh37, to call variants using VCFtools (41). GRCh37 is built from sequences from different individuals and serves as the wild type relative. If the reference allele is C at a specific genomic position, a variant is called if it is not a C. We excluded insertions and deletions from analysis. BCFtools (42) was used to slice VCFs into chromosomes (ancient DNA) and genes of interest. For 1000 Genomes, BCF tools was also used to generate population specific VCF files. BCF tools was used to create consensus sequences for each gene for the entire 1000 Genomes dataset and for each population.

We cataloged variants using two sets of VCF data published by Max Planck Institute for Evolutionary Anthropology Leipzig in 2013 and 2016. Both datasets are based on the same original sequence data but the 2016 VCFs were generated using snpAD, an ancient DNA damage-aware genotyper, and are more conservative estimates mutations (37).

The 2016 ancient VCFs (Table S2, S6) include high quality data from three Neandertals (Altai, Chagyrskaya, Vindija), lower quality data from one Neandertal (Mezmaiskaya), high quality from one Denisova (Denisovan 3), and one ancient human contemporary to Altai Neandertal and Denisova in Denisova Cave (Ust’-Ishim).

The 2013 VCF data analysis included high quality data—but not subject to the damage-aware genotyper—for Altai Neandertal, Denisova 3, and Ust’-Ishim. Variants were called using a custom bioinformations pipeline (Fig. S5). Data are found at http://cdna.eva.mpg.de/ust-ishim/VCF and http://cdna.eva.mpg.de/neandertal/altai/AltaiNeandertal/. Most variants were shared with present-day humans (Table S8). Three genes matched the human wild type (contained no variants, novel or shared) in all three samples tested (OR5K1, OR11A1, OR56A4) and 16 contained novel missense variants (Table S9). See Fig. S6 for functional results and Fig. S7 for regression results—Altai is significantly different from present-day humans and Denisova but there were few differences in terms of patterns of response.

#### Novel variant calling

R v4.1.0 (43) via R Studio v1.4.1717 (Rstudio, inc., Boston, MA, 2015) were used for analysis. Excel and csv files were read using readxl (44). The hash function in R used 1000 Genomes variant data (35) as a key to flag variants present in ancient samples but not found in present-day humans. Data were manipulated using dplyr (45) and tidyverse (46).

#### Genetic Analysis

Variants were called using DNAsp to generate Nexus files for each gene comparing a consensus sequence for each of the 26 groups in 1000 Genomes to the human reference sequence used for the published datasets (Max Planck and 1000 Genomes). Consensus sequences were used for each group in 1000 Genomes to reduce the sample size for comparison to ancient sequences which represented individuals, not groups. Biostrings (47) was used in R to create protein sequence files and call amino acid substitutions. Figures for percentage of variation by gene were generated using the R package ggplot2 with the viridis color-blind friendly palette and plotted panels using gridExtra (48). Concatenated amino acid sequences were used to infer phylogenetic relationships across populations for 30 genes (49). The phylogenetic tree was created using the R packages ape (50), seqinr (51), phylogram (52), gdsfmt (53), SNPrelate (54), and dendextend (55). Fst was calculated using gdsfmt (53) and SNPRelate (54) for Figs 1-2 and Figs S1-S2, S7- S9.

#### Primer design

ORs for extinct humans were created by mutating human ORs to match paleogenomic sequence Data using chimeric PCR. Forward and reverse PCR primers containing the desired mutation were designed to have a 56⁰C or 58⁰C annealing temperature, obtained from Integrated DNA Technologies, and diluted to 5μM.

#### Chimeric PCR

Chimeric PCR was performed using Phusion polymerase and Rho-tagged OR in a pCI vector as a template, with separate reactions using the forward primers paired with a 3’ pCI primer or reverse primers paired with a 5’ pCI primer (56). The reaction was started at 98⁰C for 30 seconds, then run for 25 cycles of the following conditions: denaturation at 98⁰C for 5 seconds, primer annealing at 55⁰C for 15 seconds, and elongation at 72⁰C for 30 seconds. The reaction was then held at 72⁰C for 5 minutes. The products resulting from the forward and reverse primers were combined for each mutant and diluted 10x with distilled water (Gibco). A second PCR was performed using Phusion polymerase, 5’ and 3’ pCI primers, and the combined and diluted products for each desired mutant as the template. The same PCR conditions were used. The products were purified using the QIAquick PCR Purification Kit (Qiagen), cut with MluI and NotI restriction enzymes, run on a 1.1% agarose gel with GelRed, and extracted using the QIAquick Gel Extraction Kit (Qiagen). The products were then ligated into Rho-tagged pCI cut with MluI and NotI using T4 ligase (New England Biolabs) and used to transform competent ampicillin-resistant E. Coli. These were plated on LB-ampicillin plates and incubated at 37⁰C overnight, then a single colony was grown in 4mL of 2XYT-ampicillin (100mg/mL) medium overnight at 37⁰C. The Denville Miniprep Purification Kit was used to lyse the bacteria and purify the plasmid DNA. The concentration of the products was determined using an Eppendorf Biophotometer, and then adjusted to 100ng/μL using TE buffer. The products were sequenced using BigDye Terminator Sequencing Kit (Applied Biosystems), purified using Sephadex (GE Healthcare), and sequenced with 3130 Genetic Analyzer (Applied Biosystems).

#### Luciferase assay

Functionality was tested by a Luciferase assay as described in Zhuang and Matsunami (16). Hana3A cells were plated in Minimum Essential Medium (MEM) containing 10% FBS (vol/vol), penicillin-treptomycin and amphotericin B on Poly-D-lysine coated 96-well plates (Corning #3843) and co-transfected using Lipofectamine2000 (Invitrogen) with the prepared Rho-tagged OR or the empty vector pCI (negative control), RTP1S, and the muscarinic receptor M3. The cells were transfected with elements from the Dual-Glo Luciferase Assay System (Promega): the cAMP response element (CRE) coupled with a luciferase reporter gene (L); and the constitutively active SV40 promoter region coupled with the Renilla luciferase (RL) reporter. In a period of 18-24 hours after transfection, the medium was replaced with 0 μM, 1 μM, 3.16 μM, 10 μM, 31.6 μM, 100 μM, or 316 μM of odorants (Sigma Aldrich) in CD293 (Gibco) containing glutamine and copper and incubated for 3.5 hours. After the addition of Dual-Glo luciferase substrate and buffer (Promega), plates were read using BMG Labtech POLARStar Optima plate reader. Promega Stop-and-Glo buffer and RL substrate were added, and plates were read again. The degree of activation was quantified for each well in Microsoft Excel using the formula (L- 400)/(RL-400) and reported on the y axis (emitted in lumens). The RL quantification is made due to variation in cell numbers in each well and the 0μM values provide the basal activity value for each OR. 0μM thus served as a comparison to identify OR response values to odorant stimulations —if the odor value on the y-axis exceeded the 0μM value, the OR was taken to have responded to the odor but if the odor value did not exceed the 0μM value, the OR was considered unresponsive. Dose responses and EC50s were determined with Graphpad Prism 7. Activity index have been calculated from dose responses by multiplying the absolute value of the logEC50 to the efficacy. If EC50 could not be determined because the dose response did not reach its plateau, logEC50 were set as the arbitrary value -2. These values were used for Table 1 along with odor descriptors and olfactory notes from Good Scents (57).

#### Flowcytometry

The cell surface expression of the Rho-tagged ORs was evaluated as described in Ikegami, de March et al. (24). HEK293 cells were plated in 35 mm plates at 25% confluency and grown overnight at 37°C and 5% CO2. The cells were then transfected with the Rho-tagged ORs and GFP using the Lipofectamine2000 reagent (ThermoFisher Scientific) in MEM supplemented with 10% FBS. 18 to 24h later, the cells were detached from the plates with cellstripper and resuspended in PBS containing 15 mM NaN3, 2% FBS in ice. The cells were then centrifuged at 4°C and resuspended and incubated for at least 30min in primary antibody (mouse anti Rho4D2, Millipore Sigma). The cells were then centrifuged at 4°C and resuspended and incubated for at least 30 min and in the dark in phycoerythrin (PE)-donkey anti-mouse F(ab’)2 Fragment antibody (Jackson Immunologicals: 715-116-150). Finally, the cells were stained with 7-Amino- actinomycin D (7AAD, Calbiochem). The cell surface expression was monitored through the PE fluorescence emitted by cells GFP positive, single, spherical, viable and 7AAD negative using a BD FACSCanto II. The results were analyzed with Flowjo. Olfr539 and Olfr541 were added to the experiment plan as positive and negative, respectively, controls of OR cell surface expression.

#### Homology model

The protocol followed a previously published method (56). 391 human OR sequences were aligned to pre-aligned sequences of 11 GPCRs including bovine rhodopsin (PDB: 1U19), human chemokine receptors CXCR4 (PDB: 3ODU) and CXCR1 (PDB: 2LNL), and human adenosine a2A receptor (PDB: 2YDV) using Jalview (58). The four experimental GPCR structures (1U19, 3ODU, 2YDV and 2LNL) were used as templates to build the human consensus OR by homology modeling with Modeller. The human consensus amino acid sequence was determined by aligning 391 human OR sequences and by selecting the most conserved amino acid for each position. Five models were obtained and the one fulfilling several constraints (binding cavity sufficiently large, no large folded structure in extra-cellular loops, all TMs folded as α-helices, a small α-helix structure between TM3 and TM4) was retained for further structural analysis.

Visualization of the models and picture generation were performed with VMD.

#### Regression Analysis

Data were analyzed in R for Windows 4.1.0, using the R Studio GUI for Windows 1.4.1717. Data were wrangled using dplyr (45) and tidyverse (46). The dose response dependent variable was fitted to distributions using fitdistrplus (59) and formed log normal distributions. The natural log was used to transform to normally distributed data for linear regression models. The car package (60) was used for ANOVA. 0μM values, as noted above, provided a control (the baseline threshold of OR activation). Because the number of cells varied across plates, the OR tested for each sample had a different 0μM. As such, removing values below the 0μM for each correlation between present-day humans and ancient DNA sample resulted in some odors missing values for one of the lineages. We compared the results of regression for testing using only active ORs (above the 0μM value) and for the full set of ORs—the results were nearly identical (Fig S8 for 2016 VCFs, Fig S9 for 2013 VCFs). Testing the full suite of responses, active or below the threshold, allowed a broader comparison of how these cells operate. Analysis was conducted on the full dataset for both between sample correlations and for mean difference testing. Plots for data visualization and results were created using ggplot2 (61) and ggpubr (62). Panels were created using cowplot (63) and ggarrange.

---

**Fig. S1.**
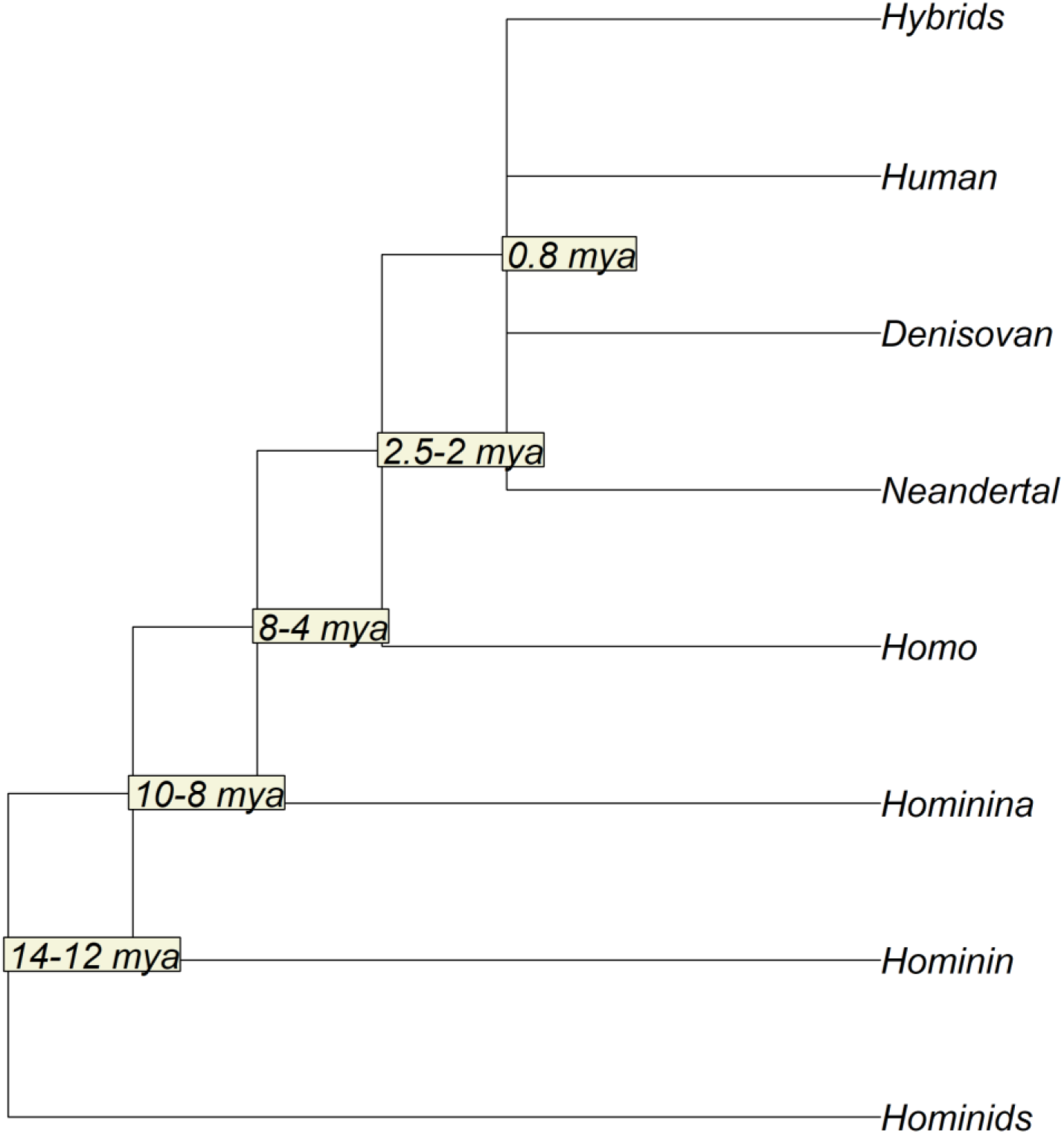
Phylogenetic tree of Hominids

**Fig. S2.**
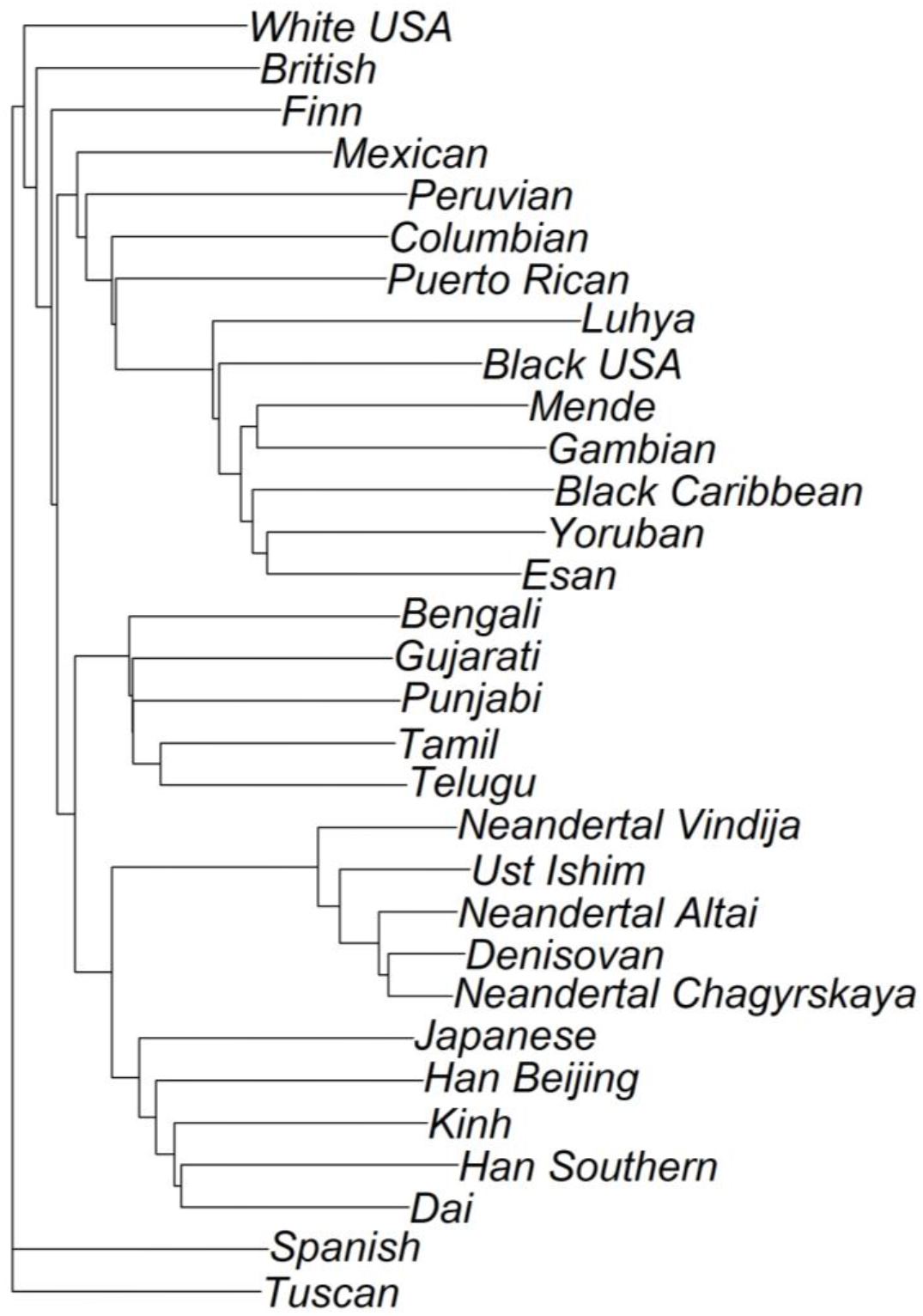
Cladogram based on full amino acid sequences for all 30 odorant receptors in ancient lineages and in 1000 Genomes populations.

**Fig. S3.**
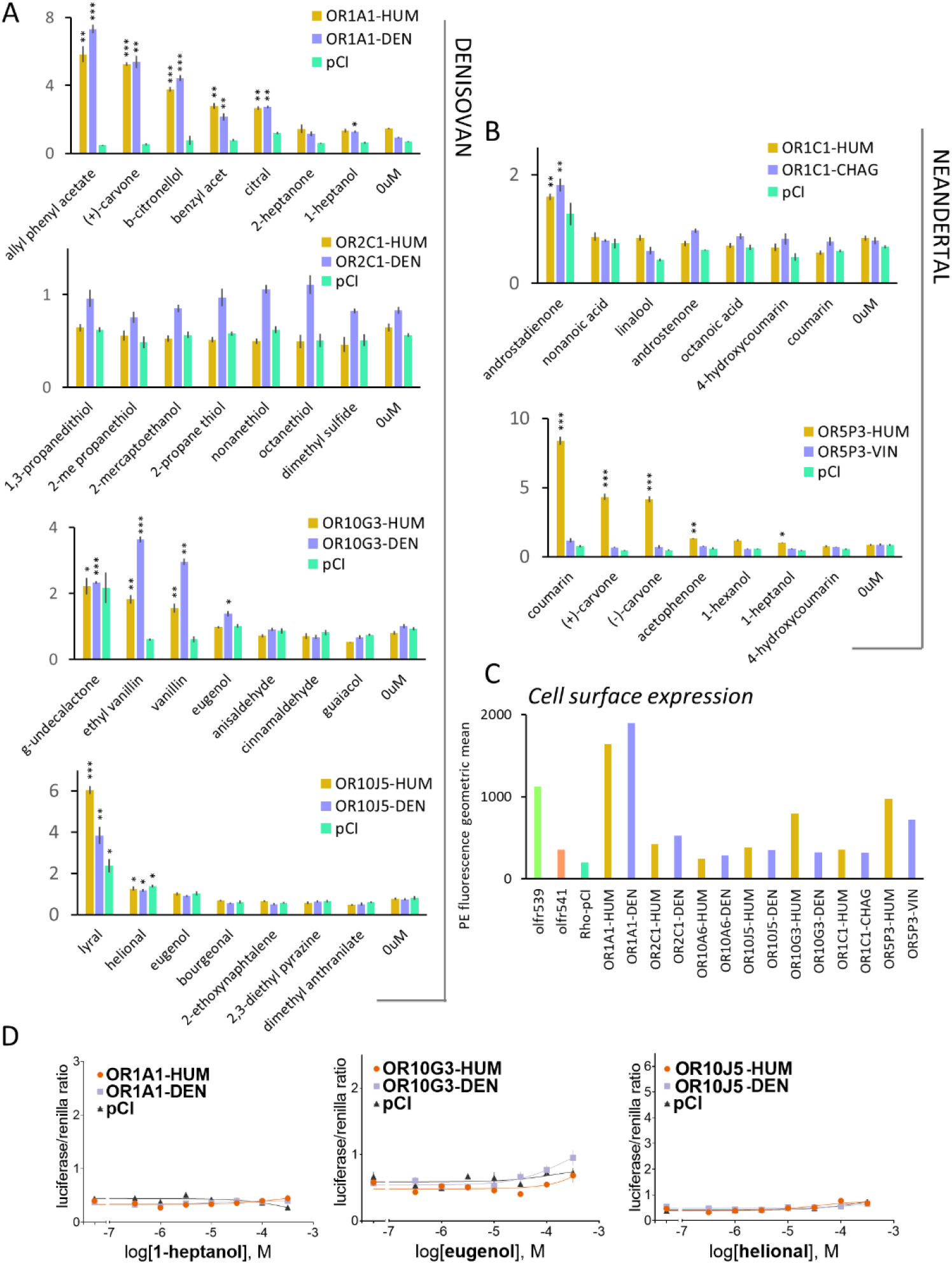
Functionality of hominin OR variants. Each Denisova (**A**) and Neandertal (**B**) OR response was first tested in luciferase assay against seven odorants (100µM) previously identified as evoking responses in human OR1A1 (21–23), OR1C1 (22), OR2C1 (21), OR10J5 (21, 22), OR5P3 (21, 22), and OR10G3 (22). The empty vector pCI was added to each experiment as a control of OR-specific response. The y-axis represents the normalized luminescence as a measure of OR activation. Asterisks represent the level of significance in a paired t-test between the odorant stimulation and the no odor (0µM) control for each OR (* p<0.05; ** p<0.01; *** p<0.001). **C**. Cell surface expression of all OR variants. Olfr539 and Olfr541, served as positive and negative control of cell surface presence, respectively. **D.** Dose response of the false positives detected in the screening (**A**).

**Fig. S4.**
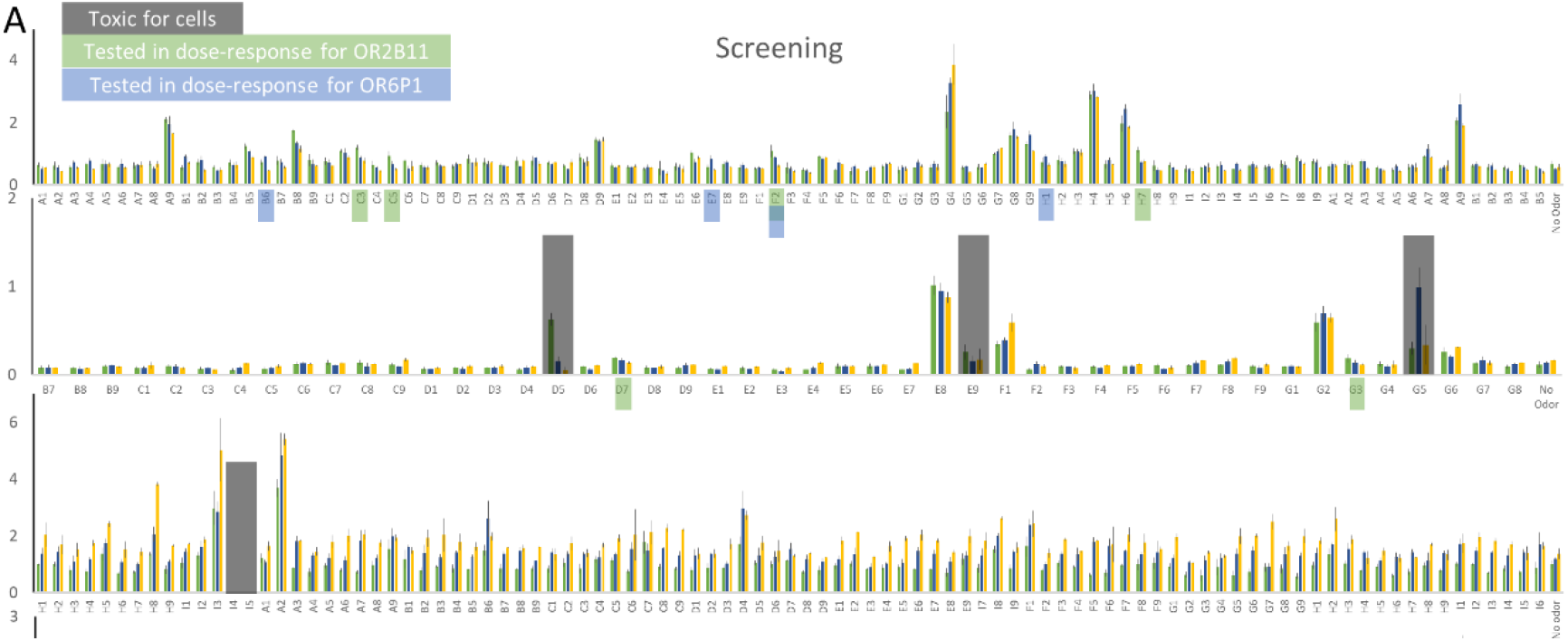
OR2B11 and OR6P1 response. **A.** Activation screening for OR2B11 and OR6P1 against more than 300 odorant molecules at 100 µM. **B**. List of odorants selected for validation by dose response. **C**. Dose-responses of OR2B11 to the odorants listed in **B**. **D.** Dose-responses of OR6P1 to the odorants listed in **B**. In panels **A**, **C** and **D** the y-axis represents the normalized luminescence response that is synonymous with OR activation.

**Fig. S5.**
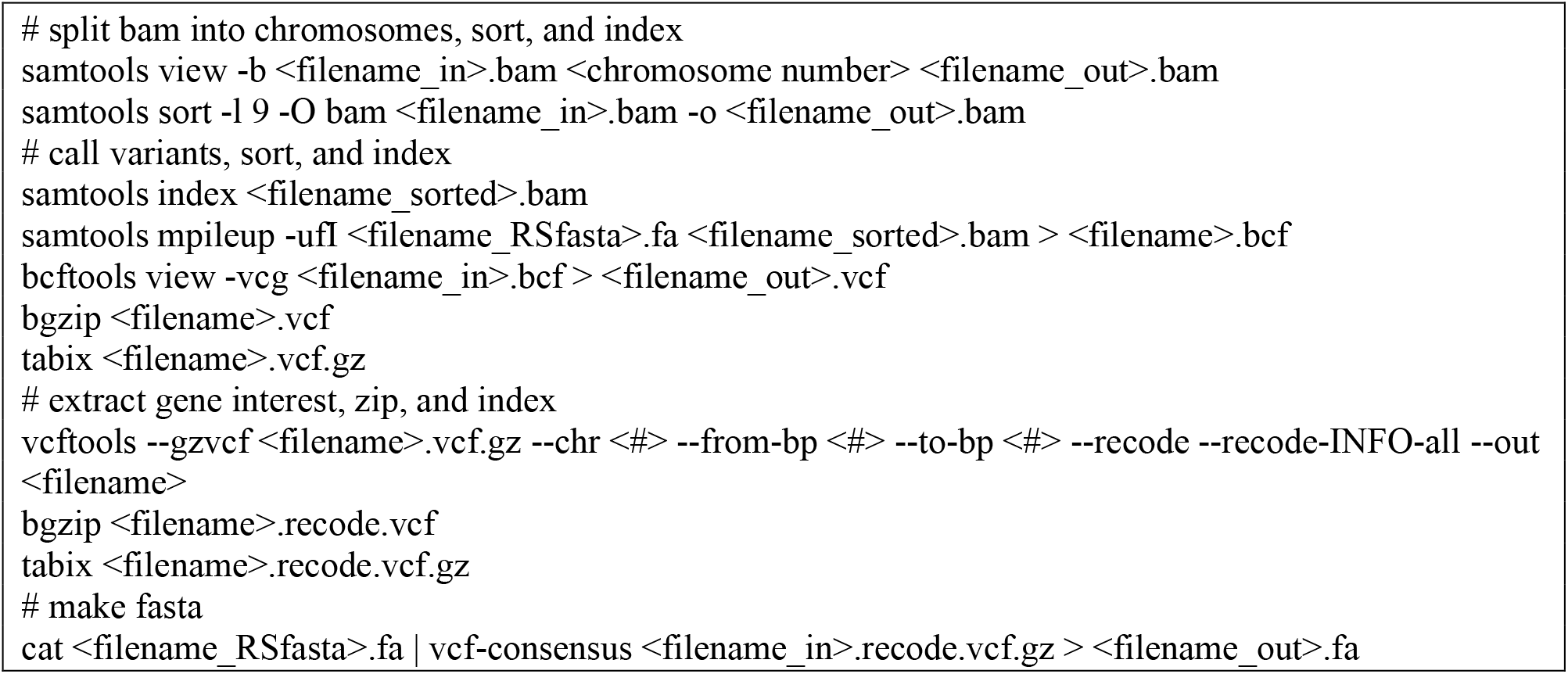
Custom variant calling pipeline for data not in VCF format

**Fig. S6.**
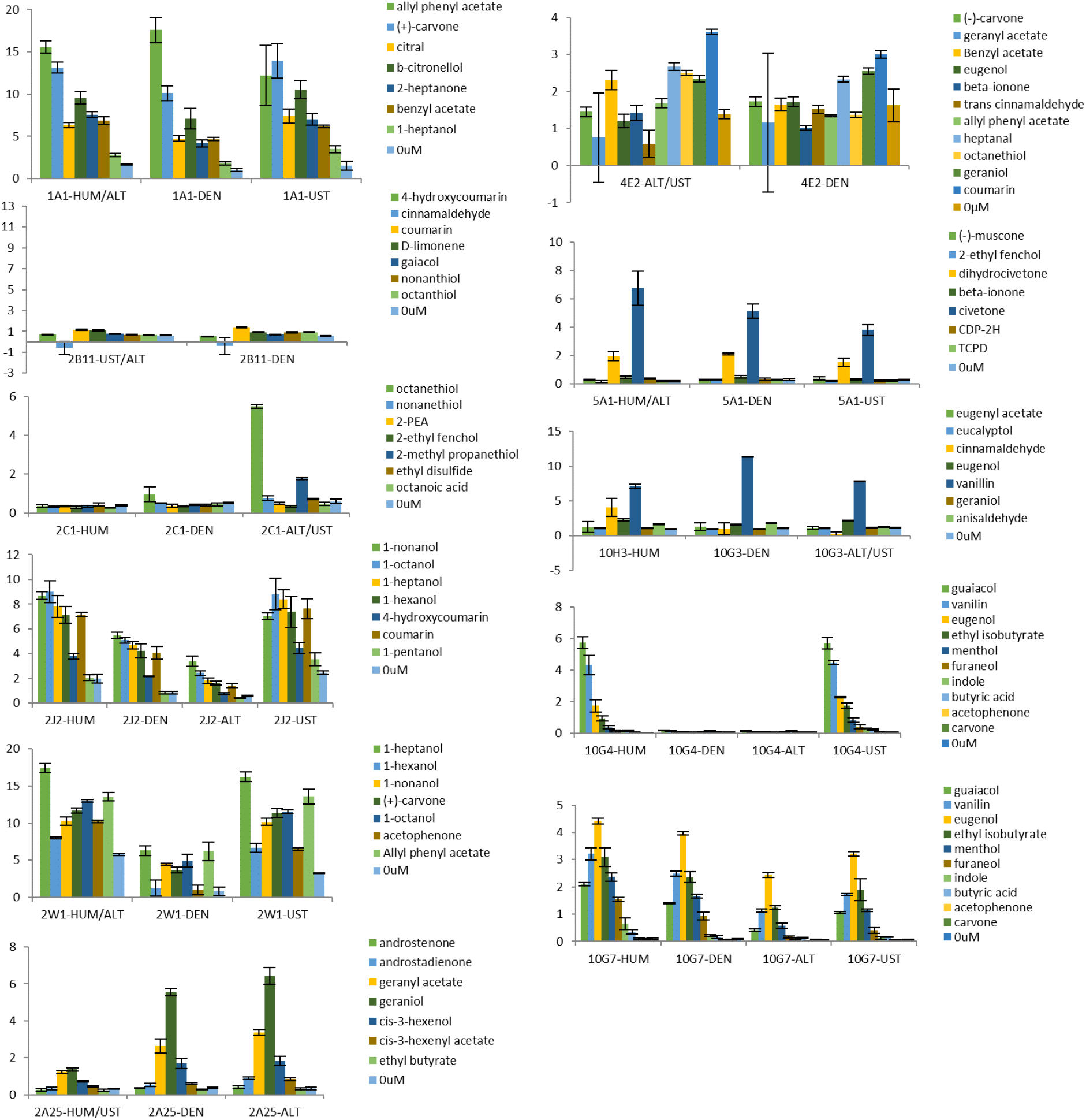
Screening of variants identified with the 2013 VCF Data and their corresponding human version response against odorants at 100µM. The y-axis shows normalized luminescence recorded in the luciferase assay.

**Fig. S7.**
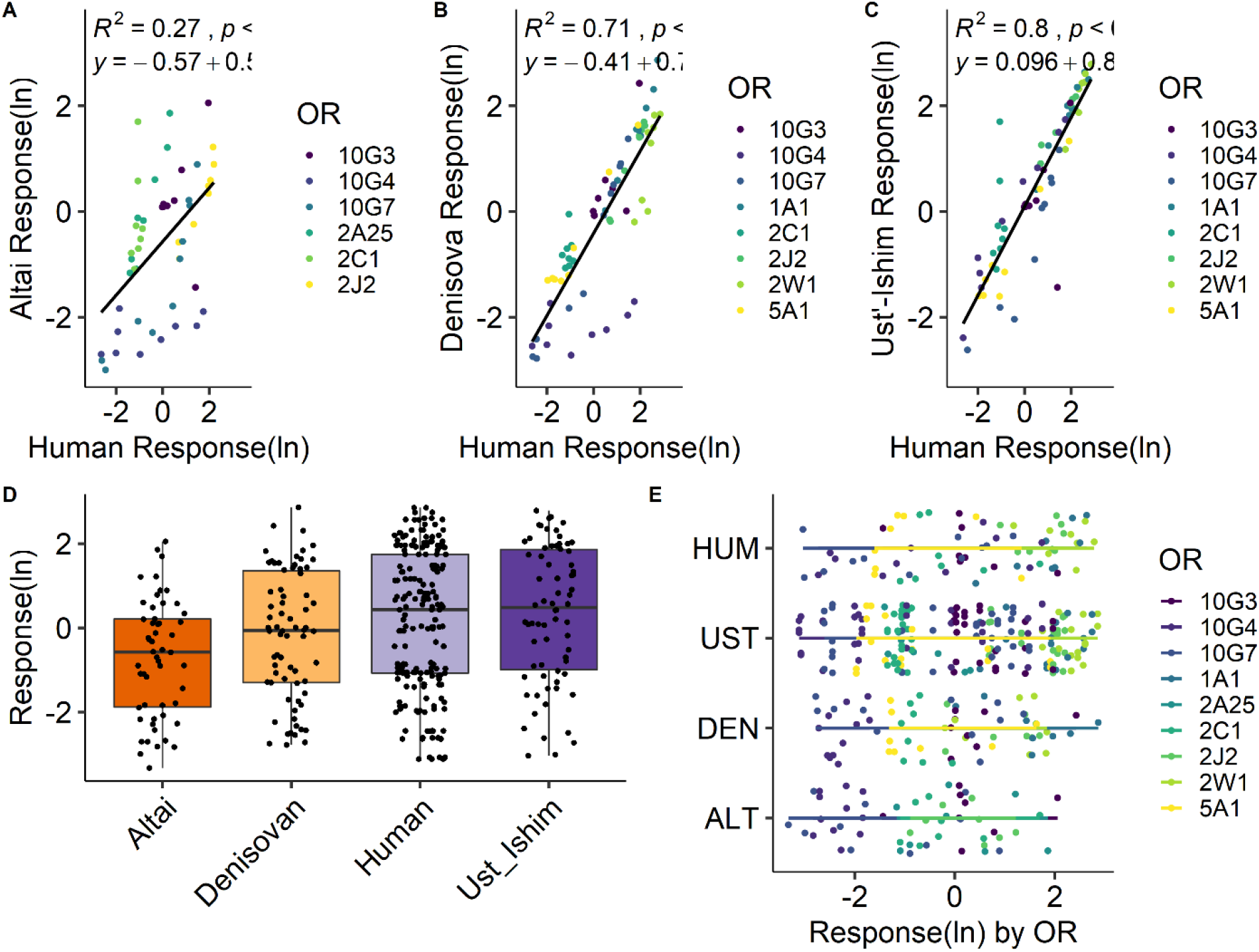
Comparison of human (x-axis) and hominin (y-axis) OR responses for **(A)** Altai Neandertal **(B)** Denisova **(C)** Ancient human hunter-gatherer, Ust’-Ishim. **D.** Boxplots of natural log of OR responses across lineages (humans are divided into ancient and modern). **E.** OR response by lineage and gene. Each OR is represented by a different color and each point represents the natural log of the response to an odorant. Dotted lines corresponds to the linear regression for the entire set of ORs responses for a given lineage. The corresponding equation and R^2^ values are shown on the regression plot.

**Fig. S8.**
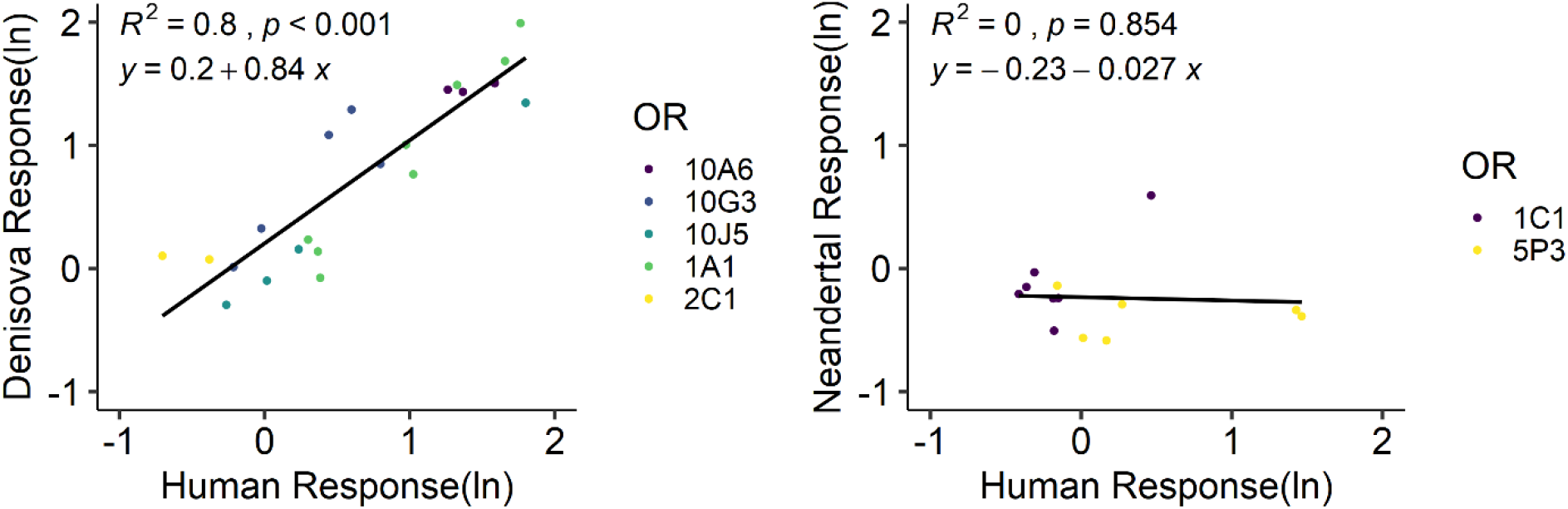
Regression results for active ORs only

**Fig. S9.**
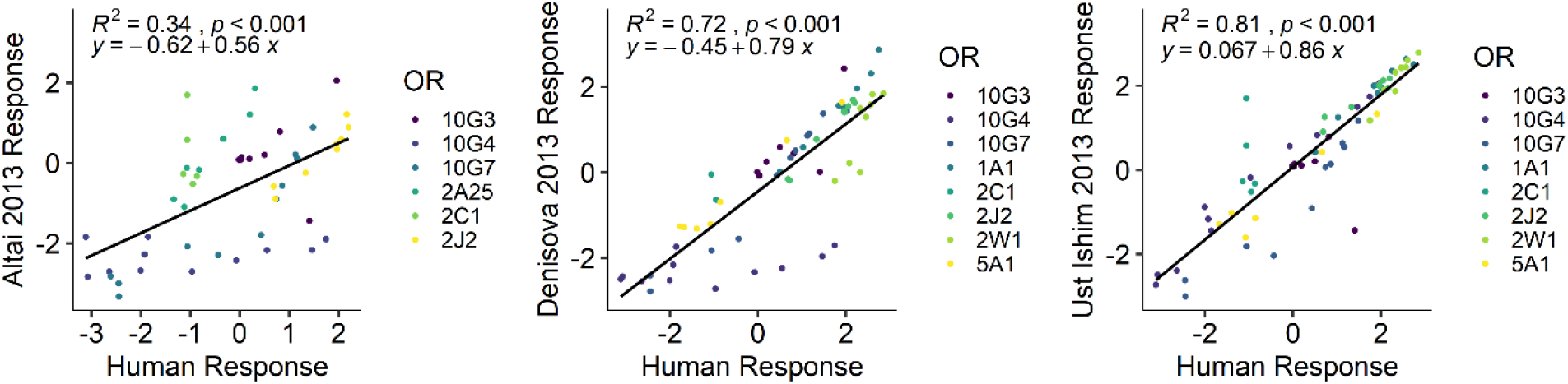
Regression results for active ORs only, 2013 data.

**Table S1.** Target Genes.

| Gene | Chr | Region | Odorant | Description |
| --- | --- | --- | --- | --- |
| OR6P1 | 1 | 158532441-158533394 | anisaldehyde | anise, licorice |
| OR10J5 | 1 | 159504868-159505797 | lyral | floral, lily |
| OR2B11 | 1 | 247614331-247615284 | coumarin | grassy, vanilla, almond |
| OR1C1 | 1 | 247920764-247921708 | linalool | floral, bergamot, lavender |
| OR5K1 | 3 | 98188421-98189347 | eugenol methyl ether | clove-like |
| OR2W1 | 6 | 29011990-29012952 | cis-3-hexen-1-ol | freshly cut grass, leaves |
| OR2J3 | 6 | 29079588-29080603 | cinnamaldehyde, cinnamon | cinnamon |
| OR2J2 | 6 | 29141413-29142351 | ethyl vanillin | vanilla |
| OR11A1 | 6 | 29394471-29395418 | 2-ethylfenchol | soil, earth, peat |
| OR2A25 | 7 | 143771313-143772245 | geranyl acetate | floral, fruity, rose |
| OR51E1 | 11 | 4673757-4674713 | isovaleric acid | sweat, cheese |
| OR51E2 | 11 | 4702979-4703941 | androsthenone, musk | β-ionone, violet-rose |
| OR51L1 | 11 | 5020213-5021160 | allyl phenyl acetate | honey |
| OR56A4 | 11 | 6023281-6024378 | undecanal | green coffee, leafy |
| OR5P3 | 11 | 7846584-7847519 | coumarin | grassy, vanilla, almond |
| OR10A6 | 11 | 7949265-7950209 | 3-phenyl propyl propionate | floral balsam hyacinth mimosa |
| OR8K3 | 11 | 56085783-56086721 | menthol | mint, menthol |
| OR5AN1 | 11 | 59131932-59132867 | muscone | musk |
| OR5A1 | 11 | 59210642-59211589 | β-ionone | violet rose |
| OR10G4 | 11 | 123886282-123887217 | vanillin | vanilla |
| OR10G7 | 11 | 123908773-123909708 | eugenol methyl ether | clove-like (fruits and spices) |
| OR8D1 | 11 | 124179736-124180662 | caramel furanone | low-burnt sugar, high-fenugreek/curry |
| OR8B3 | 11 | 124266306-124267247 | .+(-) and -(-)carvone | spearmint and caraway |
| OR4Q3 | 14 | 20215587-20216528 | eugenol methyl ether | clove-like |
| OR10G3 | 14 | 22037934-22038875 | vanillin | vanilla |
| OR4E2 | 14 | 22133297-22134238 | n-amyl acetate | bananas, apples |
| OR2C1 | 16 | 3405941-3406879 | octanethiol | sulphur |
| OR1A1 | 17 | 3118915-3119844 | -(-)carvone | spearmint |
| OR7D4 | 19 | 9324575-9325513 | androsthenone, adrostadienone | musk |
| OR7C1 | 19 | 14909986-14910948 | androstadienone | musk |

**Table S2.** Materials and Accession Numbers for Ancient Genomic Data

| Lineage | Specimen | Location | Date | Accession/DOI |
| --- | --- | --- | --- | --- |
| Denisova | Denisova3 | Denisovan<br>Cave, Altai<br>Mts,<br>Siberia | 74-82K BP | PRJEB3092<br>10.1126/science.1224344 |
| Neandertal | Altai | Denisovan<br>Cave, Altai<br>Mts,<br>Siberia | 50.3 kya ±<br>2.2 | PRJEB1265<br>10.1038/nature12886 |
|  | Chagyrskaya | Chagyrskaya<br>Cave,<br>Altai Mts,<br>Siberia | 781-126<br>kya | eva.mpg.de/neandertal/Chagyrskaya/VCF/<br>10.1016/j.aeae.2013.07.002 |
|  | Mezmaiskaya1 | Mezmaiskaya<br>Cave,<br>Russian<br>Caucasus | 60-70 kya | PRJEB21195<br>10.1126/science.aao1887 |
|  | Vindija<br>33.19 | Vindija<br>Cave,<br>Croatia | ~50 kya | PRJEB21157<br>10.1126/science.aao1887 |
| Ancient<br>Human | Ust'-Ishim | Ust'-Ishim<br>District,<br>Western<br>Siberia | 49K BP | PRJEB6622<br>10.1038/nature13810 |

In the 2016 VCF data, several variants were also found in the earlier 2013 data. A few variants in the 2013 Ust'-Ishim VCF data were not found in the 2016 dataset—due to the damaged DNA genotyper.

**Table S3.** 1000 Genomes populations

|  | **Group** | **n** |
| --- | --- | --- |
| Africa | Esan, Nigeria | 99 |
|  | Gambian | 113 |
|  | Luhya, Kenya | 99 |
|  | Mende, Sierra Leone | 85 |
|  | Yoruban, Nigeria | 108 |
| Asia | Chinese-Beijing Han | 103 |
|  | Chinese-Southern Han | 105 |
|  | Chinese-Dai | 93 |
|  | Japanese | 104 |
|  | Kinh, Vietnam | 99 |
| Southwest Asia | Bengali in Bangladesh | 86 |
|  | Gujarati, Houston | 103 |
|  | Punjabi, Pakistan | 96 |
|  | Tamil (Sri Lanka), UK | 102 |
|  | Telugu, UK | 102 |
| Europe | American White, Utah | 99 |
|  | British | 99 |
|  | Finn | 99 |
|  | Iberian, Spain | 107 |
|  | Tuscan | 107 |
| Americas | American Black | 61 |
|  | Caribbean Black | 91 |
|  | Columbian | 94 |
|  | Mexicans, LA | 64 |
|  | Peruvian | 85 |
|  | Puerto Rican | 104 |

**Table S4.**
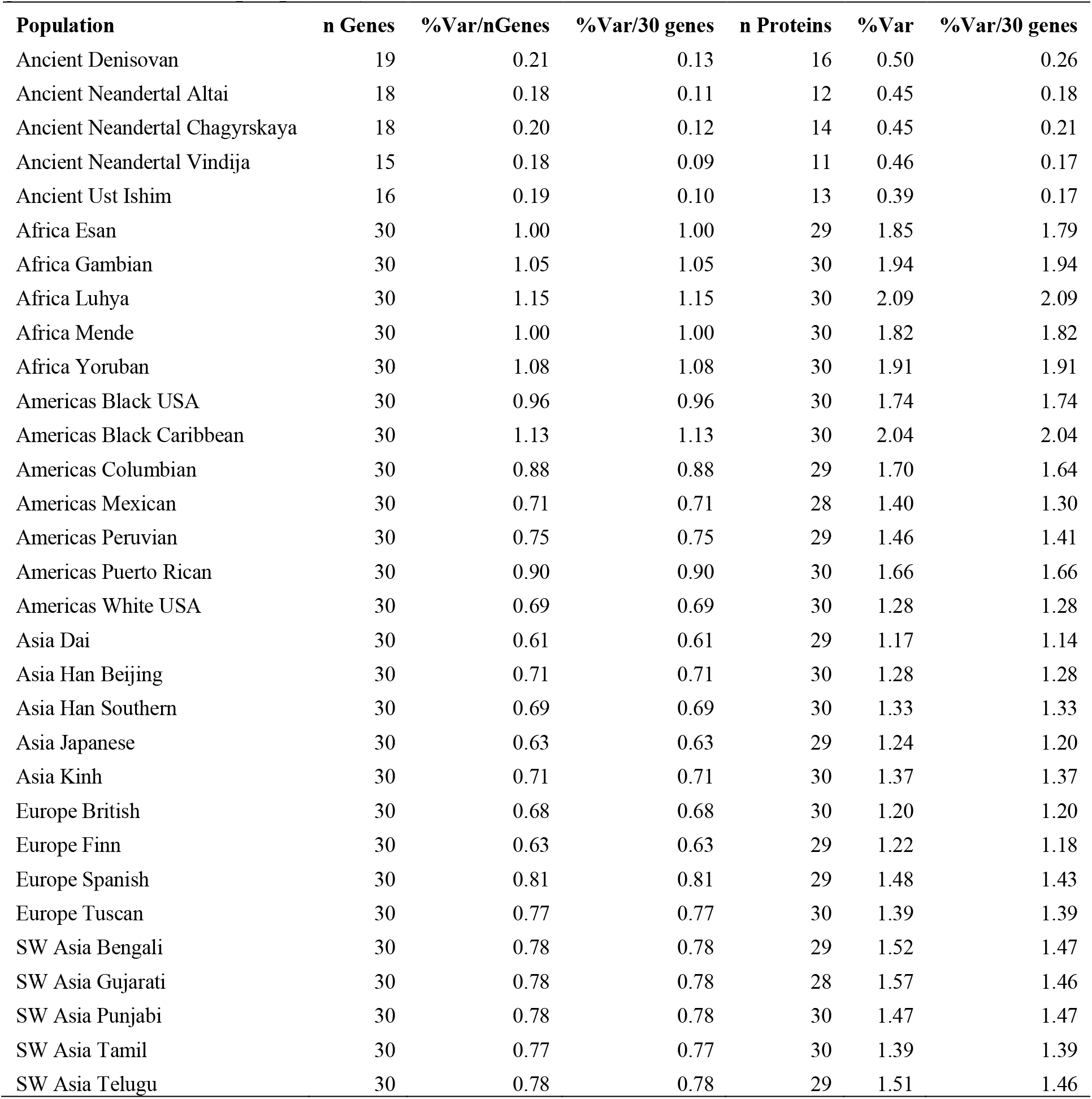
Mean of Percentage of Gene/Protein Containing Variants (e.g., n variants/total basepairs per gene or amino acids per protein)

**Table S5.**
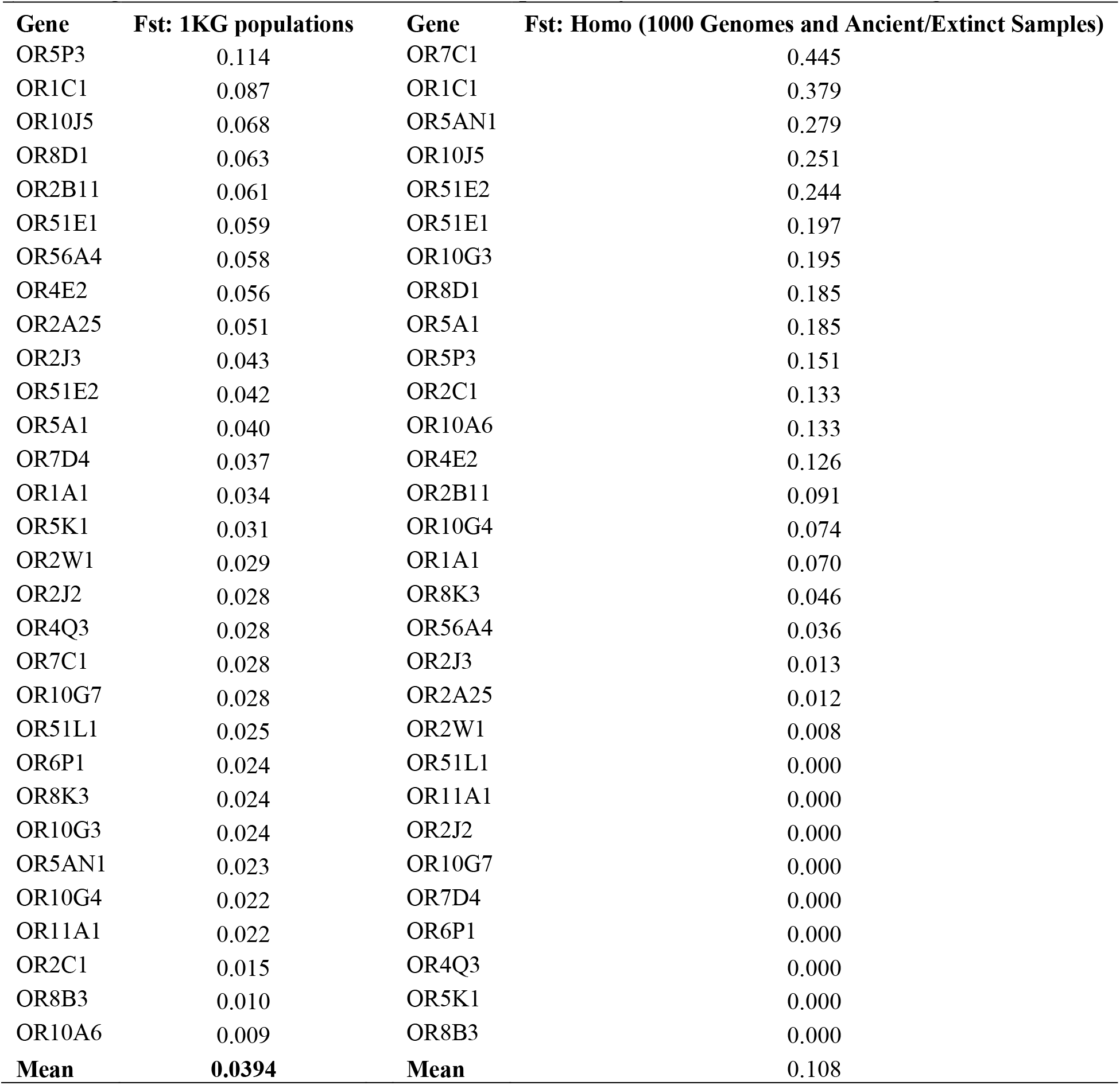
Fst based on 30 OR genes for 1000 Genomes and for all high-quality sequences for *Homo* (including 1000 Genomes. Columns are independently ordered in values from high to low.

| Gene | Fst: 1KG populations | Gene | Fst: Homo (1000 Genomes and Ancient/Extinct Samples) |
| --- | --- | --- | --- |
| OR5P3 | 0.114 | OR7C1 | 0.445 |
| OR1C1 | 0.087 | OR1C1 | 0.379 |
| OR10J5 | 0.068 | OR5AN1 | 0.279 |
| OR8D1 | 0.063 | OR10J5 | 0.251 |
| OR2B11 | 0.061 | OR51E2 | 0.244 |
| OR51E1 | 0.059 | OR51E1 | 0.197 |
| OR56A4 | 0.058 | OR10G3 | 0.195 |
| OR4E2 | 0.056 | OR8D1 | 0.185 |
| OR2A25 | 0.051 | OR5A1 | 0.185 |
| OR2J3 | 0.043 | OR5P3 | 0.151 |
| OR51E2 | 0.042 | OR2C1 | 0.133 |
| OR5A1 | 0.040 | OR10A6 | 0.133 |
| OR7D4 | 0.037 | OR4E2 | 0.126 |
| OR1A1 | 0.034 | OR2B11 | 0.091 |
| OR5K1 | 0.031 | OR10G4 | 0.074 |
| OR2W1 | 0.029 | OR1A1 | 0.070 |
| OR2J2 | 0.028 | OR8K3 | 0.046 |
| OR4Q3 | 0.028 | OR56A4 | 0.036 |
| OR7C1 | 0.028 | OR2J3 | 0.013 |
| OR10G7 | 0.028 | OR2A25 | 0.012 |
| OR51L1 | 0.025 | OR2W1 | 0.008 |
| OR6P1 | 0.024 | OR51L1 | 0.000 |
| OR8K3 | 0.024 | OR11A1 | 0.000 |
| OR10G3 | 0.024 | OR2J2 | 0.000 |
| OR5AN1 | 0.023 | OR10G7 | 0.000 |
| OR10G4 | 0.022 | OR7D4 | 0.000 |
| OR11A1 | 0.022 | OR6P1 | 0.000 |
| OR2C1 | 0.015 | OR4Q3 | 0.000 |
| OR8B3 | 0.010 | OR5K1 | 0.000 |
| OR10A6 | 0.009 | OR8B3 | 0.000 |
| **Mean** | **0.0394** | **Mean** | **0.108** |

**Table S6.**
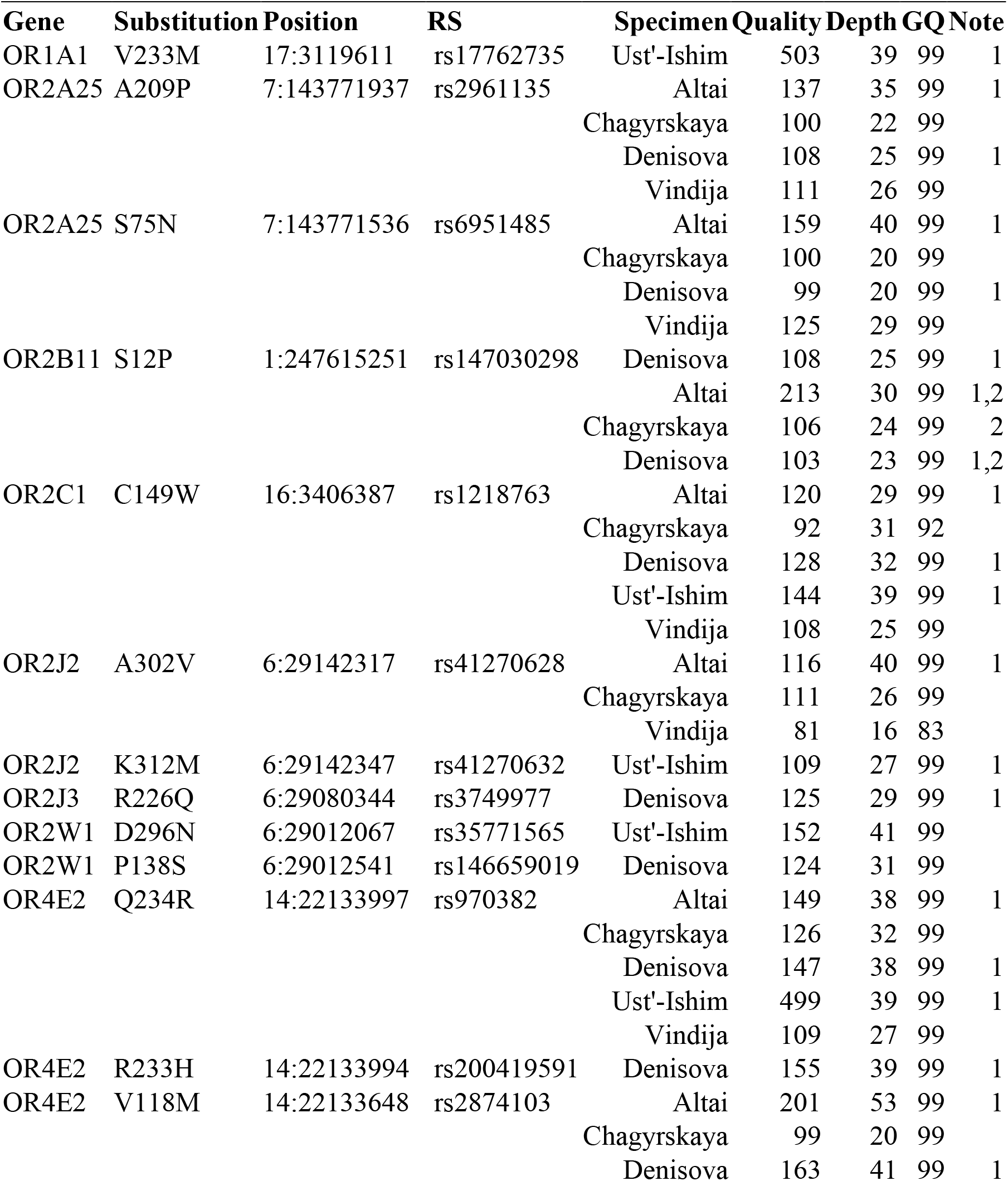
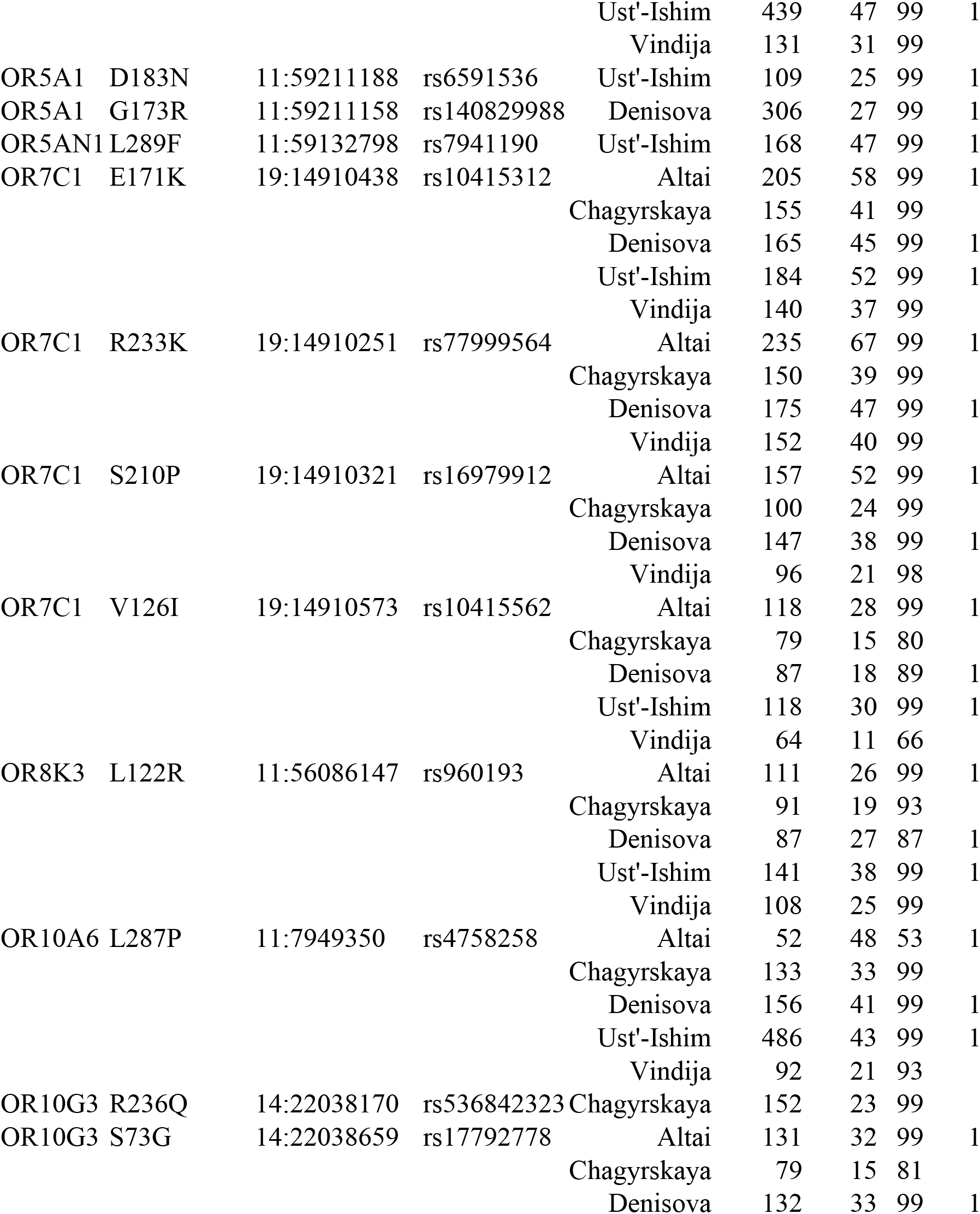
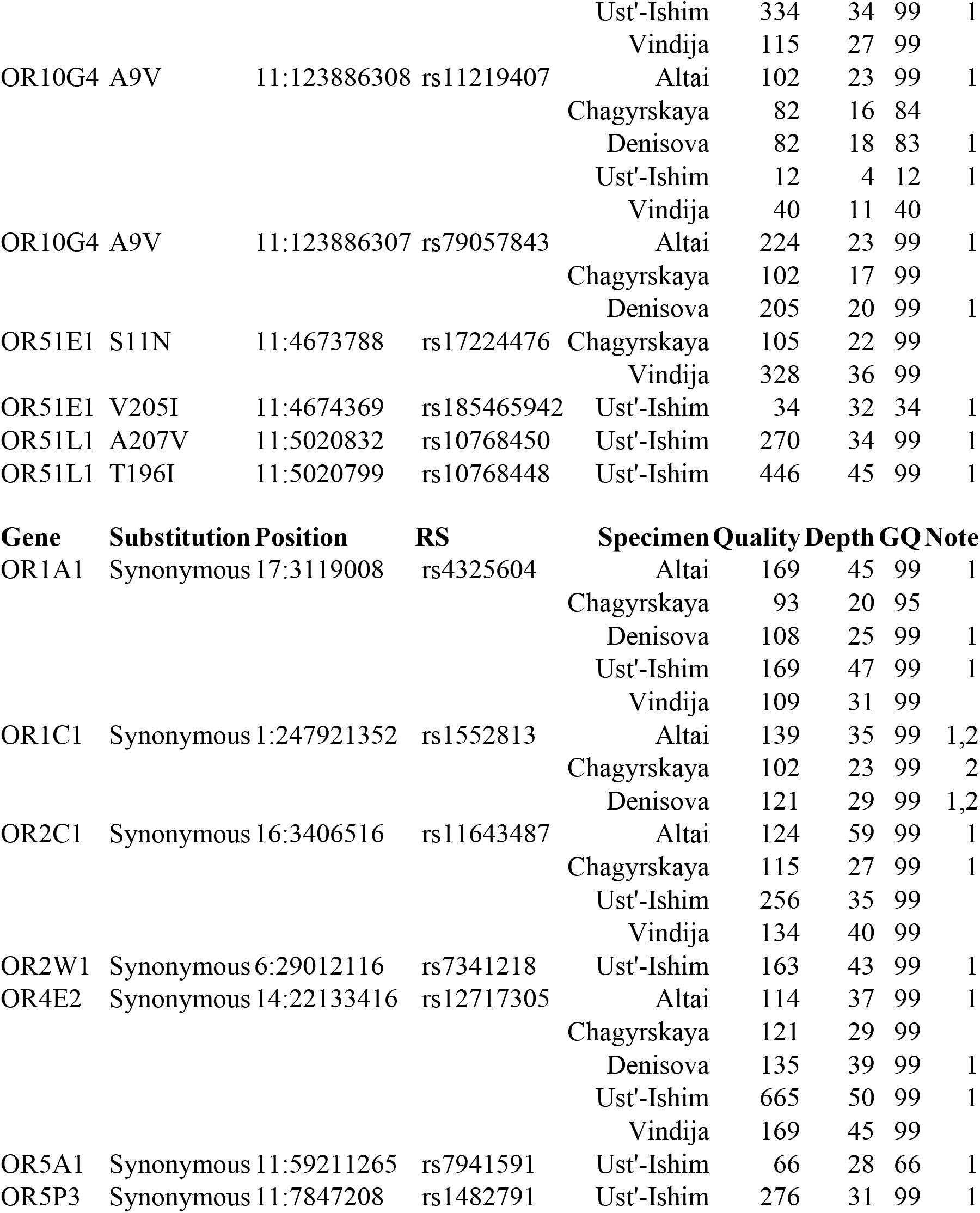
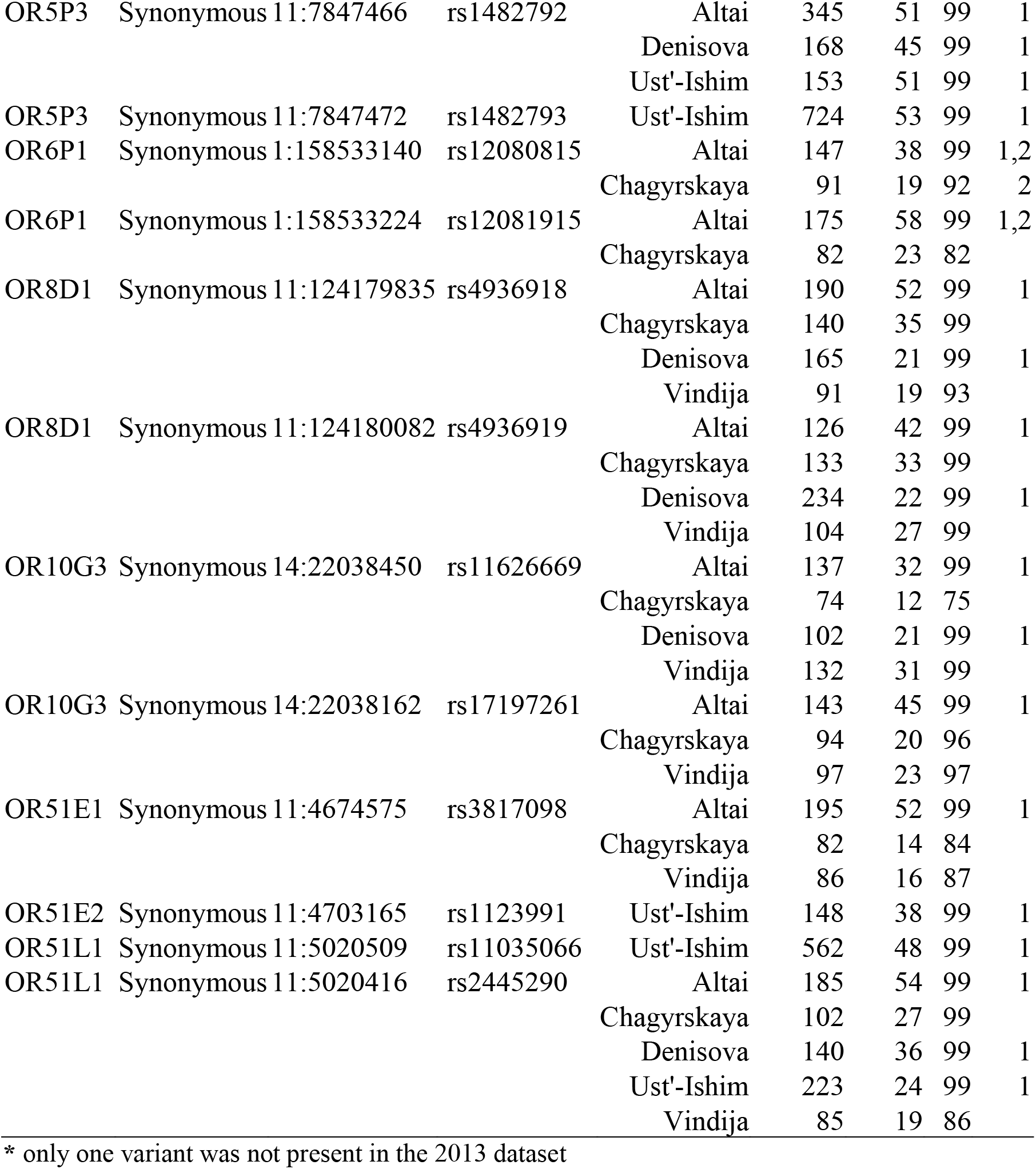
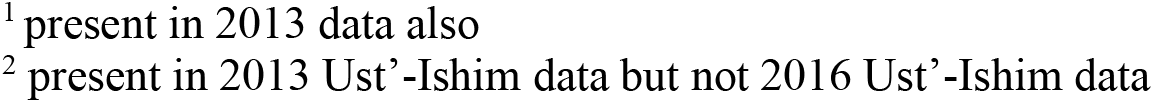
2016 VCF Shared Variants*

**Table S7.** 2016 VCF Novel Variants

| Gene | Position | Substitution | Sample | Qual | Depth | GQ | Note |
| --- | --- | --- | --- | --- | --- | --- | --- |
| OR1A1 | 17:3119050 | V46I | Denisova | 142 | 34 | 99 | 1 |
|  | 17:3119684 | T257M | Denisova | 141 | 37 | 99 | 1 |
| OR1C1 | 1:247921351 | Y120H | Chagyrskaya | 74 | 23 | 74 |  |
| OR2B11 | 1:247614839 | V149E | Denisova | 148 | 38 | 99 | 1 |
| OR2C1 | 16:3406581 | I214T | Denisova | 113 | 27 | 99 | 1 |
| OR5P3 | 11:7847045 | F159L | Vindija | 87 | 18 | 89 |  |
| OR6P1 | 1:158532749 | S216A | Altai | 174 | 47 | 99 | 1 |
|  |  |  | Chagyrskaya | 85 | 17 | 86 |  |
|  | 1:158533360 | F12S | Denisova | 102 | 23 | 99 | 1 |
| OR10G3 | 14:22038287 | E197K | Denisova | 120 | 29 | 99 | 1 |
| OR10J5 | 1:159505737 | G21R | Denisova | 142 | 48 | 99 |  |
| Gene | Position | Synonymous | Sample | Qual | Depth | GQ | Note |
| OR2B11 | 1:247615024 | V87 | Denisova | 541 | 42 | 99 | 1 |
| OR5AN1 | 11:59132645 | A238A | Chagyrskaya | 22 | 17 | 22 |  |
| OR10A6 | 11:7950075 | I45I | Denisova | 79 | 26 | 79 | 1 |
| OR10G7 | 11:123908878 | T277 | Altai | 415 | 51 | 99 | 1 |
|  |  |  | Chagyrskaya | 128 | 20 | 99 |  |
|  |  |  | Vindija | 96 | 19 | 98 |  |

<sup>1</sup> present in 2013 data also
<sup>2</sup> present in 2013 Ust'-Ishim data but not 2016 Ust'-Ishim data

**Table S8.** 2013 VCF Variants

| Specimen | Gene | Position | RS | Quality | Raw Read Depth | Mapping Quality | Type |
| --- | --- | --- | --- | --- | --- | --- | --- |
| Altai | OR6P1 | 1:158532749 | Novel | 228 | 48 | 37 | S216A |
| Altai | OR6P1 | 1:158533140 | rs12080815 | 228 | 42 | 39 | SYNON |
| Ust'-Ishim | OR6P1 | 1:158533140 | rs12080815 | 228 | 33 | 36 | SYNON |
| Altai | OR6P1 | 1:158533224 | rs12081915 | 228 | 62 | 37 | SYNON |
| Ust'-Ishim | OR6P1 | 1:158533224 | rs12081915 | 228 | 41 | 37 | SYNON |
| Denisova | OR6P1 | 1:158533360 | Novel | 228 | 20 | 37 | F12S |
| Ust'-Ishim | OR10J5 | 1:159505297 | rs4656837 | 222 | 41 | 37 | SYNON |
| Denisova | OR10J5 | 1:159505737 | Novel | 228 | 44 | 37 | G21R |
| Altai | OR2B11 | 1:247614693 | rs6695302 | 161 | 33 | 38 | V198M |
| Denisova | OR2B11 | 1:247614693 | rs6695302 | 228 | 23 | 37 | V198M |
| Ust'-Ishim | OR2B11 | 1:247614693 | rs6695302 | 228 | 34 | 37 | V198M |
| Denisova | OR2B11 | 1:247614839 | Novel | 228 | 37 | 37 | V149E |
| Altai | OR2B11 | 1:247614896 | rs11583410 | 228 | 30 | 37 | I130S |
| Denisova | OR2B11 | 1:247614896 | rs11583410 | 228 | 25 | 37 | I130S |
| Ust'-Ishim | OR2B11 | 1:247614896 | rs11583410 | 228 | 20 | 37 | I130S |
| Denisova | OR2B11 | 1:247615024 | Novel | 222 | 39 | 37 | SYNON |
| Denisova | OR2B11 | 1:247615251 | rs147030298 | 228 | 25 | 37 | S12P |
| Ust'-Ishim | OR1C1 | 1:247921100 | rs1552812 | 228 | 36 | 37 | SYNON |
| Altai | OR1C1 | 1:247921352 | rs1552813 | 228 | 37 | 38 | SYNON |
| Denisova | OR1C1 | 1:247921352 | rs1552813 | 228 | 29 | 37 | SYNON |
| Ust'-Ishim | OR1C1 | 1:247921352 | rs1552813 | 228 | 33 | 37 | SYNON |
| Ust'-Ishim | OR5K1 | 3:98188365 | rs9861575 | 182 | 50 | 37 | W14C |
| Ust'-Ishim | OR2W1 | 6:29012067 | rs35771565 | 228 | 42 | 38 | D296N |
| Ust'-Ishim | OR2W1 | 6:29012116 | rs7341218 | 228 | 44 | 38 | SYNON |
| Denisova | OR2W1 | 6:29012541 | rs146659019 | 228 | 31 | 37 | P138A |
| Denisova | OR2J3 | 6:29080344 | rs3749977 | 228 | 27 | 37 | R226Q |
| Altai | OR2J3 | 6:29080349 | rs3130764 | 228 | 36 | 36 | I228V |
| Denisova | OR2J3 | 6:29080349 | rs3130764 | 228 | 26 | 35 | I228V |
| Altai | OR2J3 | 6:29080450 | rs3130765 | 228 | 44 | 31 | M261I/M |
| Denisova | OR2J3 | 6:29080450 | rs3130765 | 228 | 35 | 26 | M261I/M |
| Altai | OR2J2 | 6:29141772 | rs3129158 | 228 | 50 | 31 | SYNON |
| Denisova | OR2J2 | 6:29141772 | rs3129158 | 203 | 20 | 28 | SYNON |
| Altai | OR2J2 | 6:29142317 | rs41270628 | 228 | 44 | 37 | A302V |
| Ust'-Ishim | OR2J2 | 6:29142347 | rs41270632 | 228 | 31 | 34 | K312M |
| Altai | OR2A25 | 7:143771536 | rs6951485 | 228 | 45 | 35 | S75N |
| Denisova | OR2A25 | 7:143771536 | rs6951485 | 228 | 22 | 33 | S75N |
| Denisova | OR2A25 | 7:143771733 | Novel | 136 | 32 | 37 | I141V |
| Altai | OR2A25 | 7:143771937 | rs2961135 | 228 | 43 | 38 | A209P |
| Denisova | OR2A25 | 7:143771937 | rs2961135 | 228 | 25 | 37 | A209P |
| Ust'-Ishim | OR51E1 | 11:4674369 | rs185465942 | 33 | 32 | 37 | V205I |
| Altai | OR51E1 | 11:4674575 | rs3817098 | 228 | 56 | 37 | SYNON |
| Ust'-Ishim | OR51E2 | 11:4703165 | rs1123991 | 228 | 39 | 37 | SYNON |
| Altai | OR51L1 | 11:5020416 | rs2445290 | 228 | 60 | 38 | SYNON |
| Denisova | OR51L1 | 11:5020416 | rs2445290 | 228 | 36 | 37 | SYNON |
| Ust'-Ishim | OR51L1 | 11:5020416 | rs2445290 | 156 | 24 | 37 | SYNON |
| Ust'-Ishim | OR51L1 | 11:5020509 | rs11035066 | 222 | 52 | 36 | SYNON |
| Ust'-Ishim | OR51L1 | 11:5020799 | rs10768448 | 222 | 45 | 36 | T196I |
| Ust'-Ishim | OR51L1 | 11:5020832 | rs10768450 | 222 | 37 | 37 | A207V |
| Altai | OR5P3 | 11:7846722 | Novel | 222 | 46 | 37 | Y266STOP |
| Altai | OR5P3 | 11:7846752 | Novel | 222 | 52 | 37 | SYNON |
| Ust'-Ishim | OR5P3 | 11:7847208 | rs1482791 | 157 | 32 | 37 | SYNON |
| Altai | OR5P3 | 11:7847466 | rs1482792 | 222 | 55 | 39 | SYNON |
| Denisova | OR5P3 | 11:7847466 | rs1482792 | 228 | 44 | 37 | SYNON |
| Ust'-Ishim | OR5P3 | 11:7847466 | rs1482792 | 228 | 54 | 36 | SYNON |
| Ust'-Ishim | OR5P3 | 11:7847472 | rs1482793 | 222 | 56 | 36 | SYNON |
| Altai | OR10A6 | 11:7949350 | rs4758258 | 228 | 54 | 39 | SYNON |
| Denisova | OR10A6 | 11:7949350 | rs4758258 | 228 | 42 | 37 | SYNON |
| Ust'-Ishim | OR10A6 | 11:7949350 | rs4758258 | 222 | 45 | 38 | SYNON |
| Ust'-Ishim | OR10A6 | 11:7950024 | rs12272735 | 183 | 28 | 34 | SYNON |
| Denisova | OR10A6 | 11:7950075 | Novel | 228 | 26 | 37 | SYNON |
| Altai | OR8K3 | 11:56086147 | rs960193 | 228 | 29 | 38 | L122R |
| Denisova | OR8K3 | 11:56086147 | rs960193 | 228 | 24 | 35 | L122R |
| Ust'-Ishim | OR8K3 | 11:56086147 | rs960193 | 228 | 41 | 36 | L122R |
| Ust'-Ishim | OR5AN1 | 11:59132798 | rs7941190 | 228 | 50 | 37 | L289F |
| Denisova | OR5A1 | 11:59211158 | rs140829988 | 222 | 29 | 37 | G173R |
| Ust'-Ishim | OR5A1 | 11:59211188 | rs6591536 | 228 | 25 | 37 | D183N |
| Ust'-Ishim | OR5A1 | 11:59211265 | rs7941591 | 228 | 28 | 37 | SYNON |
| Altai | OR10G4 | 11:123886307 | rs79057843 | 175 | 43 | 28 | A9V |
| Denisova | OR10G4 | 11:123886307 | rs79057843 | 160 | 25 | 32 | A9V |
| Altai | OR10G4 | 11:123886308 | rs11219407 | 228 | 43 | 28 | A9V |
| Denisova | OR10G4 | 11:123886308 | rs11219407 | 228 | 25 | 32 | A9V |
| Ust'-Ishim | OR10G4 | 11:123886308 | rs11219407 | 89 | 11 | 21 | A9V |
| Ust'-Ishim | OR10G4 | 11:123886320 | Novel | 88 | 7 | 19 | T13A |
| Altai | OR10G4 | 11:123886681 | Novel | 193 | 59 | 14 | M134V |
| Denisova | OR10G4 | 11:123886681 | Novel | 44 | 33 | 10 | M134V |
| Altai | OR10G4 | 11:123886865 | rs4084209 | 228 | 31 | 33 | V195E |
| Denisova | OR10G4 | 11:123886865 | rs4084209 | 228 | 19 | 30 | V195E |
| Ust'-Ishim | OR10G4 | 11:123886865 | rs4084209 | 228 | 21 | 28 | V195E |
| Denisova | OR10G4 | 11:123886948 | Novel | 50 | 32 | 17 | S223T |
| Denisova | OR10G4 | 11:123886950 | Novel | 76 | 32 | 17 | S223T |
| Altai | OR10G4 | 11:123886984 | rs4936880 | 221 | 50 | 34 | R235G |
| Denisova | OR10G4 | 11:123886984 | rs4936880 | 228 | 19 | 36 | R235G |
| Altai | OR10G4 | 11:123887164 | rs4936881 | 228 | 32 | 25 | K295Q |
| Denisova | OR10G4 | 11:123887164 | rs4936881 | 205 | 17 | 22 | K295Q |
| Altai | OR10G4 | 11:123887184 | rs4936882 | 228 | 28 | 33 | SYNON |
| Denisova | OR10G4 | 11:123887184 | rs4936882 | 228 | 23 | 34 | SYNON |
| Ust'-Ishim | OR10G4 | 11:123887184 | rs4936882 | 228 | 33 | 33 | SYNON |
| Altai | OR10G7 | 11:123908878 | Novel | 222 | 55 | 37 | SYNON |
| Altai | OR10G7 | 11:123908977 | rs188316662 | 143 | 54 | 15 | SYNON |
| Denisova | OR10G7 | 11:123909073 | rs11219419 | 29 | 30 | 23 | SYNON |
| Ust'-Ishim | OR10G7 | 11:123909073 | rs11219419 | 228 | 25 | 27 | SYNON |
| Altai | OR10G7 | 11:123909099 | rs59358830 | 222 | 37 | 37 | L204I |
| Ust'-Ishim | OR10G7 | 11:123909172 | rs28414940 | 74 | 43 | 8 | SYNON |
| Altai | OR10G7 | 11:123909302 | rs513591 | 175 | 41 | 33 | T136S |
| Denisova | OR10G7 | 11:123909302 | rs513591 | 150 | 17 | 34 | T136S |
| Altai | OR10G7 | 11:123909311 | rs472442 | 75 | 32 | 30 | N133S |
| Denisova | OR10G7 | 11:123909311 | rs472442 | 126 | 15 | 34 | N133S |
| Ust'-Ishim | OR10G7 | 11:123909342 | rs470242 | 200 | 12 | 27 | SYNON |
| Altai | OR10G7 | 11:123909637 | Novel | 70 | 8 | 34 | SYNON |
| Ust'-Ishim | OR10G7 | 11:123909650 | rs3894199 | 111 | 32 | 17 | G20A |
| Altai | OR10G7 | 11:123909653 | rs4998340 | 29 | 17 | 22 | P19Q |
| Ust'-Ishim | OR10G7 | 11:123909653 | rs4998340 | 135 | 31 | 18 | P19Q |
| Altai | OR10G7 | 11:123909656 | rs3894198 | 20 | 21 | 21 | A18L |
| Denisova | OR10G7 | 11:123909656 | rs3894198 | 10 | 7 | 19 | A18L |
| Ust'-Ishim | OR10G7 | 11:123909656 | rs3894198 | 128 | 28 | 17 | A18L |
| Altai | OR10G7 | 11:123909671 | rs11827843 | 113 | 27 | 23 | T13M |
| Denisova | OR10G7 | 11:123909671 | rs11827843 | 47 | 16 | 20 | T13M |
| Altai | OR10G7 | 11:123909695 | rs3894197 | 203 | 38 | 35 | T5S |
| Denisova | OR10G7 | 11:123909695 | rs3894197 | 87 | 15 | 26 | T5S |
| Ust'-Ishim | OR10G7 | 11:123909695 | rs3894197 | 228 | 24 | 29 | T5S |
| Altai | OR8D1 | 11:124179835 | rs4936918 | 228 | 53 | 37 | SYNON |
| Denisova | OR8D1 | 11:124179835 | rs4936918 | 126 | 19 | 37 | SYNON |
| Altai | OR8D1 | 11:124180082 | rs4936919 | 228 | 45 | 38 | SYNON |
| Denisova | OR8D1 | 11:124180082 | rs4936919 | 193 | 20 | 37 | SYNON |
| Ust'-Ishim | OR8B3 | 11:124266906 | Novel | 221 | 64 | 26 | SYNON |
| Altai | OR8B3 | 11:124266912 | rs28398895 | 228 | 55 | 33 | SYNON |
| Denisova | OR8B3 | 11:124266912 | rs28398895 | 228 | 34 | 34 | SYNON |
| Ust'-Ishim | OR8B3 | 11:124266912 | rs28398895 | 220 | 61 | 27 | SYNON |
| Altai | OR8B3 | 11:124267056 | rs144478570 | 38 | 61 | 12 | SYNON |
| Denisova | OR8B3 | 11:124267056 | rs144478570 | 15 | 63 | 12 | SYNON |
| Denisova | OR8B3 | 11:124267189 | rs507335 | 10 | 9 | 2 | H20R |
| Altai | OR4Q3 | 14:20216298 | rs12896533 <sup>1</sup> | 228 | 35 | 29 | F238L |
| Denisova | OR4Q3 | 14:20216298 | rs12896533 <sup>1</sup> | 228 | 24 | 28 | F238L |
| Ust'-Ishim | OR4Q3 | 14:20216298 | rs12896533 <sup>1</sup> | 228 | 33 | 27 | F238L |
| Altai | OR10G3 | 14:22038162 | rs17197261 | 228 | 51 | 38 | SYNON |
| Denisova | OR10G3 | 14:22038287 | Novel | 228 | 27 | 37 | SYNON |
| Altai | OR10G3 | 14:22038450 | rs11626669 | 228 | 35 | 39 | SYNON |
| Denisova | OR10G3 | 14:22038450 | rs11626669 | 228 | 17 | 37 | SYNON |
| Ust'-Ishim | OR10G3 | 14:22038525 | rs11626693 | 222 | 50 | 37 | SYNON |
| Altai | OR10G3 | 14:22038659 | rs17792778 | 228 | 37 | 39 | S73G |
| Denisova | OR10G3 | 14:22038659 | rs17792778 | 228 | 29 | 37 | S73G |
| Ust'-Ishim | OR10G3 | 14:22038659 | rs17792778 | 222 | 36 | 36 | S73G |
| Altai | OR4E2 | 14:22133416 | rs12717305 | 228 | 40 | 38 | SYNON |
| Denisova | OR4E2 | 14:22133416 | rs12717305 | 228 | 38 | 37 | SYNON |
| Ust'-Ishim | OR4E2 | 14:22133416 | rs12717305 | 222 | 51 | 37 | SYNON |
| Altai | OR4E2 | 14:22133648 | rs2874103 | 228 | 57 | 39 | V118M |
| Denisova | OR4E2 | 14:22133648 | rs2874103 | 228 | 40 | 37 | V118M |
| Ust'-Ishim | OR4E2 | 14:22133648 | rs2874103 | 222 | 47 | 37 | V118M |
| Denisova | OR4E2 | 14:22133994 | rs200419591 | 228 | 29 | 37 | R233H |
| Altai | OR4E2 | 14:22133997 | rs970382 | 228 | 39 | 37 | Q234R |
| Denisova | OR4E2 | 14:22133997 | rs970382 | 228 | 31 | 37 | Q234R |
| Ust'-Ishim | OR4E2 | 14:22133997 | rs970382 | 222 | 39 | 37 | Q234R |
| Altai | OR2C1 | 16:3405994 | rs1228348 | 228 | 62 | 38 | SYNON |
| Ust'-Ishim | OR2C1 | 16:3405994 | rs1228348 | 165 | 27 | 37 | SYNON |
| Altai | OR2C1 | 16:3406387 | rs1218763 | 228 | 33 | 37 | C149W |
| Denisova | OR2C1 | 16:3406387 | rs1218763 | 228 | 31 | 37 | C149W |
| Ust'-Ishim | OR2C1 | 16:3406387 | rs1218763 | 228 | 41 | 37 | C149W |
| Altai | OR2C1 | 16:3406516 | rs11643487 | 228 | 59 | 37 | SYNON |
| Ust'-Ishim | OR2C1 | 16:3406516 | rs11643487 | 146 | 36 | 37 | SYNON |
| Denisova | OR2C1 | 16:3406581 | Novel | 228 | 29 | 37 | I214T |
| Altai | OR1A1 | 17:3119008 | rs4325604 | 228 | 46 | 36 | SYNON |
| Denisova | OR1A1 | 17:3119008 | rs4325604 | 228 | 19 | 37 | SYNON |
| Ust'-Ishim | OR1A1 | 17:3119008 | rs4325604 | 228 | 48 | 37 | SYNON |
| Denisova | OR1A1 | 17:3119050 | Novel | 228 | 30 | 37 | V46I |
| Denisova | OR1A1 | 17:3119383 | Novel | 228 | 40 | 37 | L157F |
| Altai | OR1A1 | 17:3119397 | rs769425 | 228 | 59 | 37 | SYNON |
| Ust'-Ishim | OR1A1 | 17:3119397 | rs769425 | 222 | 39 | 35 | SYNON |
| Ust'-Ishim | OR1A1 | 17:3119611 | rs17762735 | 222 | 40 | 37 | V233M |
| Denisova | OR1A1 | 17:3119684 | Novel | 228 | 27 | 37 | T257M |
| Denisova | OR7D4 | 19:9324904 | rs546470301 | 221 | 35 | 26 | A204T |
| Denisova | OR7D4 | 19:9324971 | Novel | 139 | 36 | 31 | SYNON |
| Altai | OR7C1 | 19:14910251 | rs77999564 | 228 | 66 | 37 | R233K |
| Denisova | OR7C1 | 19:14910251 | rs77999564 | 228 | 43 | 37 | R233K |
| Altai | OR7C1 | 19:14910321 | rs16979912 | 228 | 55 | 37 | S210P |
| Denisova | OR7C1 | 19:14910321 | rs16979912 | 228 | 38 | 37 | S210P |
| Altai | OR7C1 | 19:14910438 | rs10415312 | 228 | 58 | 37 | E171K |
| Denisova | OR7C1 | 19:14910438 | rs10415312 | 228 | 36 | 37 | E171K |
| Ust'-Ishim | OR7C1 | 19:14910438 | rs10415312 | 228 | 52 | 37 | E171K |
| Altai | OR7C1 | 19:14910573 | rs10415562 | 228 | 36 | 36 | V126I |
| Denisova | OR7C1 | 19:14910573 | rs10415562 | 228 | 24 | 33 | V126I |
| Ust'-Ishim | OR7C1 | 19:14910573 | rs10415562 | 228 | 44 | 31 | V126I |

<sup>1</sup> rs12896533 is no longer considered valid (AKA rs61086986)

**Table S9.** Novel 2013 VCF Variants

| **Specimen** | **Gene** | **Position** | **RS** | **Quality** | **Raw**<br>**Read**<br>**Depth** | **Mapping**<br>**Quality** | **Type** |
| --- | --- | --- | --- | --- | --- | --- | --- |
| Altai | OR10G4 | 11:123886681 | Novel | 193 | 59 | 14 | M134V |
| Altai | OR10G7 | 11:123908878 | Novel | 222 | 55 | 37 | SYNON |
| Altai | OR10G7 | 11:123909637 | Novel | 70 | 8 | 34 | SYNON |
| Altai | OR5P3 | 11:7846722 | Novel | 222 | 46 | 37 | Y266STOP |
| Altai | OR5P3 | 11:7846752 | Novel | 222 | 52 | 37 | SYNON |
| Altai | OR6P1 | 1:158532749 | Novel | 228 | 48 | 37 | S216A |
| Denisova | OR10A6 | 11:7950075 | Novel | 228 | 26 | 37 | SYNON |
| Denisova | OR10G3 | 14:22038287 | Novel | 228 | 27 | 37 | SYNON |
| Denisova | OR10G4 | 11:123886681 | Novel | 44 | 33 | 10 | M134V |
| Denisova | OR10G4 | 11:123886948 | Novel | 50 | 32 | 17 | S223T |
| Denisova | OR10G4 | 11:123886950 | Novel | 76 | 32 | 17 | S223T |
| Denisova | OR10J5 | 1:159505737 | Novel | 228 | 44 | 37 | G21R |
| Denisova | OR1A1 | 17:3119050 | Novel | 228 | 30 | 37 | V46I |
| Denisova | OR1A1 | 17:3119383 | Novel | 228 | 40 | 37 | L157F |
| Denisova | OR1A1 | 17:3119684 | Novel | 228 | 27 | 37 | T257M |
| Denisova | OR2A25 | 7:143771733 | Novel | 136 | 32 | 37 | I141V |
| Denisova | OR2B11 | 1:247614839 | Novel | 228 | 37 | 37 | V149E |
| Denisova | OR2B11 | 1:247615024 | Novel | 222 | 39 | 37 | SYNON |
| Denisova | OR2C1 | 16:3406581 | Novel | 228 | 29 | 37 | I214T |
| Denisova | OR6P1 | 1:158533360 | Novel | 228 | 20 | 37 | F12S |
| Denisova | OR7D4 | 19:9324971 | Novel | 139 | 36 | 31 | SYNON |
| Ust'-Ishim | OR10G4 | 11:123886320 | Novel | 88 | 7 | 19 | T13A |
| Ust'-Ishim | OR8B3 | 11:124266906 | Novel | 221 | 64 | 26 | SYNON |

## Data S1. (separate file)

Raw data and codes for this project and data derived from it are available in at https://github.com/kchoover14/OldNoses and are usable under the license provided in the repository.

## Notes

### Competing Interest Statement

The authors have declared no competing interest.

### Summary of Updates

This version has a revised Fst table and the data that underlie that calculation. There were some minor edits to text following Fst revision and first review process.

## References and Notes

1. G. M. Hughes et al., The birth and death of olfactory receptor gene families in mammalian niche adaptation. Mol. Biol. Evol., msy028–msy028 (2018).

2. S. Hayden et al., Ecological adaptation determines functional mammalian olfactory subgenomes. Genome Res. 20, 1–9 (2010).

3. A. Varki, D. H. Geschwind, E. E. Eichler, Human uniqueness: genome interactions with environment, behaviour and culture. Nature Reviews Genetics 9, 749 (2008).

4. I. C. Winder et al., Evolution and dispersal of the genus Homo: A landscape approach. J Hum Evol 87, 48–65 (2015).

5. K. Prufer et al., The complete genome sequence of a Neanderthal from the Altai Mountains. Nature 505, 43–49 (2014).

6. Z. Jacobs et al., Timing of archaic hominin occupation of Denisova Cave in southern Siberia. Nature 565, 594–599 (2019).

7. E. I. Zavala et al., Pleistocene sediment DNA reveals hominin and faunal turnovers at Denisova Cave. Nature, (2021).

8. F. Chen et al., A late Middle Pleistocene Denisovan mandible from the Tibetan Plateau. Nature 569, 409–412 (2019).

9. G. S. Jacobs et al., Multiple Deeply Divergent Denisovan Ancestries in Papuans. Cell 177, 1010–1021.e1032 (2019).

10. F. Luca, G. H. Perry, D. R. A., Evolutionary Adaptations to Dietary Changes. Annu. Rev. Nutr. 30, 291–314 (2010).

11. Jeremy F. McRae et al., Identification of Regions Associated with Variation in Sensitivity to Food-Related Odors in the Human Genome. Curr. Biol. 23, 1596–1600 (2013).

12. S. R. Jaeger et al., A Mendelian trait for olfactory sensitivity affects odor experience and food selection. Curr. Biol. 23, 1601–1605 (2013).

13. J. F. McRae et al., Genetic Variation in the Odorant Receptor OR2J3 Is Associated with the Ability to Detect the “Grassy” Smelling Odor, cis-3-hexen-1-ol. Chem. Senses 37, 585–593 (2012).

14. K. Lunde et al., Genetic variation of an odorant receptor OR7D4 and sensory perception of cooked meat containing androstenone. PLoS One 7, e35259 (2012).

15. K. C. Hoover et al., Global survey of variation in a human olfactory receptor gene reveals signatures of non-neutral evolution. Chem. Senses 40, 481–488 (2015).

16. H. Zhuang, M. Chien, H. Matsunami, Dynamic functional evolution of an odorant receptor for sex-steroid-derived odors in primates. Proc Natl Acad Sci U S A 106, 21247–21251 (2009).

17. J. D. Mainland et al., The missense of smell: functional variability in the human odorant receptor repertoire. Nat Neurosci 17, 114–120 (2014).

18. F. Mafessoni et al., A high-coverage Neandertal genome from Chagyrskaya Cave. Proceedings of the National Academy of Sciences 117, 15132–15136 (2020).

19. S. Wright, The interpretation of population structure by f-statistics with special regard to systems of mating. Evolution 19, 395–420 (1965).

20. A. R. Templeton, Human Races: A Genetic and Evolutionary Perspective. American Anthropologist 100, 632–650 (1998).

21. H. Saito, Q. Chi, H. Zhuang, H. Matsunami, J. Mainland, Odor coding by a Mammalian receptor repertoire. Sci Signal 2, ra9 (2009).

22. K. A. Adipietro, J. D. Mainland, H. Matsunami, Functional evolution of mammalian odorant receptors. PLoS Genet. 8, e1002821 (2012).

23. K. Schmiedeberg et al., Structural determinants of odorant recognition by the human olfactory receptors OR1A1 and OR1A2. J. Struct. Biol. 159, 400–412 (2007).

24. K. Ikegami et al., Structural instability and divergence from conserved residues underlie intracellular retention of mammalian odorant receptors. Proc Natl Acad Sci U S A 117, 2957–2967 (2020).

25. M. Bastir et al., Evolution of the base of the brain in highly encephalized human species. Nature Communications 2, 588 (2011).

26. T. Weiss et al., Human Olfaction without Apparent Olfactory Bulbs. Neuron, (2019).

27. F. W. Marlowe, et al., Honey, Hadza, hunter-gatherers, and human evolution. J. Hum. Evol. 71, 119–128 (2014).

28. N. J. Dominy, Ferment in the family tree. Proceedings of the National Academy of Sciences 112, 308–309 (2015).

29. J. A. Fellows Yates et al., The evolution and changing ecology of the African hominid oral microbiome. Proceedings of the National Academy of Sciences 118, e2021655118 (2021).

30. G. Wang, J. E. Hayes, G. R. Ziegler, R. F. Roberts, H. Hopfer, Dose-Response Relationships for Vanilla Flavor and Sucrose in Skim Milk: Evidence of Synergy. Beverages 4, 73 (2018).

31. Y. Gilad, O. Man, G. Glusman, A comparison of the human and chimpanzee olfactory receptor gene repertoires. Genome Research 15, 224–230 (2005).

32. M. Somel et al., A scan for human-specific relaxation of negative selection reveals unexpected polymorphism in proteasome genes. Mol. Biol. Evol. 30, 1808–1815 (2013).

33. J. Lachance et al., Evolutionary history and adaptation from high-coverage whole- genome sequences of diverse African hunter-gatherers. Cell 150, 457–469 (2012).

34. C. Trimmer et al., Genetic variation across the human olfactory receptor repertoire alters odor perception. Proceedings of the National Academy of Sciences, 201804106 (2019).

35. The Genomes Project Consortium, The 1000 Genomes Project Consortium. Nature 526, 68–74 (2015).

36. G. M. Hughes, E. C. Teeling, D. G. Higgins, Loss of olfactory receptor function in hominin evolution. PLOS ONE 9, e84714 (2014).

37. K. Prüfer et al., A high-coverage Neandertal genome from Vindija Cave in Croatia. Science, (2017).

38. F. Mafessoni, et al., A high-coverage Neandertal genome from Chagyrskaya Cave. bioRxiv, (2020).

39. L. L. Prieto-Godino, et al., Olfactory receptor pseudo-pseudogenes. Nature 539, 93–97 (2016).

40. I. H. Barnes et al., Expert curation of the human and mouse olfactory receptor gene repertoires identifies conserved coding regions split across two exons. BMC Genomics 21, 1–15 (2020).

41. P. Danecek et al., The variant call format and VCFtools. Bioinformatics 27, 2156–2158 (2011).

42. P. Danecek, S. A. McCarthy, BCFtools/csq: haplotype-aware variant consequences. Bioinformatics 33, 2037–2039 (2017).

43. R. C. Team. (2021).

44. H. Wickham et al. (2019).

45. H. F. Wickham, Romain; Henry, Lionel; Müller, Kirill, in R package version 1.0.7. (2021).

46. H. Wickham, The tidyverse. R package ver 1, 836 (2017).

47. H. Pagès, P. Aboyoun, R. Gentleman, S. DebRoy, Biostrings: Efficient manipulation of biological strings. R package version 2, 10.18129 (2019).

48. B. Auguie, A. Antonov, M. B. Auguie, Package ‘gridExtra’. Miscellaneous Functions for “Grid” Graphics, (2017).

49. S. R. Gadagkar, M. S. Rosenberg, S. Kumar, Inferring species phylogenies from multiple genes: concatenated sequence tree versus consensus gene tree. Journal of Experimental Zoology Part B: Molecular and Developmental Evolution 304, 64–74 (2005).

50. E. Paradis, J. Claude, K. Strimmer, APE: analyses of phylogenetics and evolution in R language. Bioinformatics 20, 289–290 (2004).

51. D. Charif, J. R. Lobry, in Structural approaches to sequence evolution. (Springer, 2007), pp. 207–232.

52. S. P. Wilkinson, S. K. Davy, phylogram: an R package for phylogenetic analysis with nested lists. Journal of Open Source Software 3, 790 (2018).

53. X. Zheng, M. X. Zheng, Package ‘gdsfmt’. (2014).

54. X. Zheng, M. X. Zheng, Package ‘SNPRelate’. A package for Parallel Computing Toolset for Relatedness and Principal Component Analysis of SNP Data, (2013).

55. T. Galili, dendextend: an R package for visualizing, adjusting and comparing trees of hierarchical clustering. Bioinformatics 31, 3718–3720 (2015).

56. C. Bushdid, C. A. de March, H. Matsunami, J. Golebiowski, in Olfactory Receptors: Methods and Protocols, F. M. Simoes de Souza, G. Antunes, Eds. (Springer New York, New York, NY, 2018), pp. 77–93.

57. TheGoodScentsCompany. (2021), vol. 2021.

58. C. A. de March, S.-K. Kim, S. Antonczak, W. A. Goddard, J. Golebiowski, G protein- coupled odorant receptors: From sequence to structure. Protein Science 24, 1543–1548 (2015).

59. M. L. Delignette-Muller, C. Dutang, fitdistrplus: An R package for fitting distributions. Journal of statistical software 64, 1–34 (2015).

60. J. Fox, et al., Package ‘car’. Vienna: R Foundation for Statistical Computing, 16 (2012).

61. H. Wickham, ggplot2. Wiley Interdisciplinary Reviews: Computational Statistics 3, 180–185 (2011).

62. A. Kassambara, M. A. Kassambara. (2020).

63. C. O. Wilke, H. Wickham, M. C. O. Wilke, Package ‘cowplot’. Streamlined Plot Theme and Plot Annotations for ‘ggplot2, (2019).

